# MTHFD1 is a genetic interactor of BRD4 and links folate metabolism to transcriptional regulation

**DOI:** 10.1101/439422

**Authors:** Sara Sdelci, André F. Rendeiro, Philipp Rathert, Gerald Hofstätter, Anna Ringler, Herwig P. Moll, Wanhui You, Kristaps Klavins, Bettina Gürtl, Matthias Farlik, Sandra Schick, Freya Klepsch, Matthew Oldach, Pisanu Buphamalai, Fiorella Schischlik, Peter Májek, Katja Parapatics, Christian Schmidl, Michael Schuster, Thomas Penz, Dennis L. Buckley, Otto Hudecz, Richard Imre, Robert Kralovics, Keiryn L. Bennett, Andre C. Müller, Karl Mechtler, Jörg Menche, James E. Bradner, Georg E. Winter, Emilio Casanova, Christoph Bock, Johannes Zuber, Stefan Kubicek

## Abstract

The histone acetyl-reader BRD4 is an important regulator of chromatin structure and transcription, yet factors modulating its activity have remained elusive. Here we describe two complementary screens for genetic and physical interactors of BRD4, which converge on the folate pathway enzyme MTHFD1. We show that a fraction of MTHFD1 resides in the nucleus, where it is recruited to distinct genomic loci by direct interaction with BRD4. Inhibition of either BRD4 or MTHFD1 results in similar changes in nuclear metabolite composition and gene expression, and pharmacologic inhibitors of the two pathways synergize to impair cancer cell viability *in vitro* and *in vivo*. Our finding that MTHFD1 and other metabolic enzymes are chromatin-associated suggests a direct role for nuclear metabolism in the control of gene expression.

---

BRD4 is an important chromatin regulator with roles in gene regulation, DNA damage, cell proliferation and cancer progression^1–4^. The protein is recruited to distinct genomic loci by the interaction of its tandem bromodomains with acetylated lysines on histones and other nuclear proteins^5^. There, BRD4 acts as a transcriptional activator by P-TEFb-mediated stimulation of transcriptional elongation^6^. The activating function of BRD4 on key driver oncogenes like MYC have made this epigenetic enzyme an important therapeutic target in both BRD4 translocated and BRD4 wild-type cancers^3,7–12^, and at least seven bromodomain inhibitors have reached the clinical stage^13^. Genome-wide studies have identified the role of BRD4-induced epigenetic heterogeneity in cancer cell resistance^14^, and factors defining BRD4 inhibitor response^15,16^. However, despite its clinical importance and the broad role of BRD4 in chromatin organization, surprisingly little is known about factors that are directly required for BRD4 function. To systematically expand the list of known BRD4 interactors^5^ and to characterize proteins directly required for BRD4 function, we developed a strategy of two complementary screens for genetic and physical partners of BRD4. The two approaches converge on a single factor, methylenetetrahydrofolate dehydrogenase 1 (MTHFD1). Our description of a transcriptional role for this C-1-tetrahydrofolate synthase highlights a direct connection between nuclear folate metabolism and cancer gene regulation.

We recently generated reporter for epigenetic drug screening (REDS) cell lines that respond to inhibition of BRD4 with the expression of red fluorescent protein (RFP) (Extended Data Fig. 1a)^17^. The near-haploid genotype of these cells, originating from the chronic myeloid leukemia cell line KBM7, makes them ideally suited for gene-trap genetic screens^18,19^. We therefore tested several REDS clones for their karyotype and confirmed the haploidity of REDS1 (Extended Data Fig. 1b-c). This clone, which harbors a single RFP integration in the first intron of *CDKAL1* (Extended Data Fig. 1d-e), robustly induced RFP expression when treated with the BRD4 inhibitor (*S*)-JQ1 or with short hairpin RNAs (shRNAs) against the protein (Extended Data Fig. 1f-g). We then performed a gene-trap-mediated genetic screen in order to identify new functional interactors of BRD4 (Fig. 1a). The high specificity of the screening system relies on a rapid gain of RFP signal, which indicates chromatin changes phenocopying BRD4 inhibition. Therefore, the expression of RFP upon a specific gene knock-out suggests that the gene targeted is either directly required for BRD4 function or independently involved in chromatin remodeling at BRD4-dependent loci. We infected REDS1 cells with a gene-trap virus carrying a green fluorescent protein (GFP) reporter gene and after two weeks sorted double positive cells (RFP^+^/GFP^+^) (Fig. 1b). Following the extraction of genomic DNA from this population, gene-trap integration sites were amplified, sequenced and mapped onto the genome. Two prominent protein-coding genes emerged from the analysis of these data for the number and orientation of integrations: *MTHFD1* and *MDC1* (mediator of DNA damage checkpoint 1) (Fig. 1c, Extended Data Fig. 2a and Extended Data Table 1). *MDC1*, a gene involved in DNA repair, can indirectly be linked to BRD4 biology through the insulator role of the short isoform of BRD4 during DNA damage signaling^2^. To validate *MTHFD1* as a genetic interactor of BRD4, REDS1 cells were treated with three different shRNAs resulting in 44-92% knock-down of MTHFD1 (Fig. 1d). All three hairpins induced RFP expression, and the effect size correlated with their knock-down efficiency (Fig. 1e-f). To rule out clone-specific effects, we repeated the same experiment in the diploid REDS3 clone and obtained comparable results (Extended Data Fig. 2b-d).

**Figure 1.**
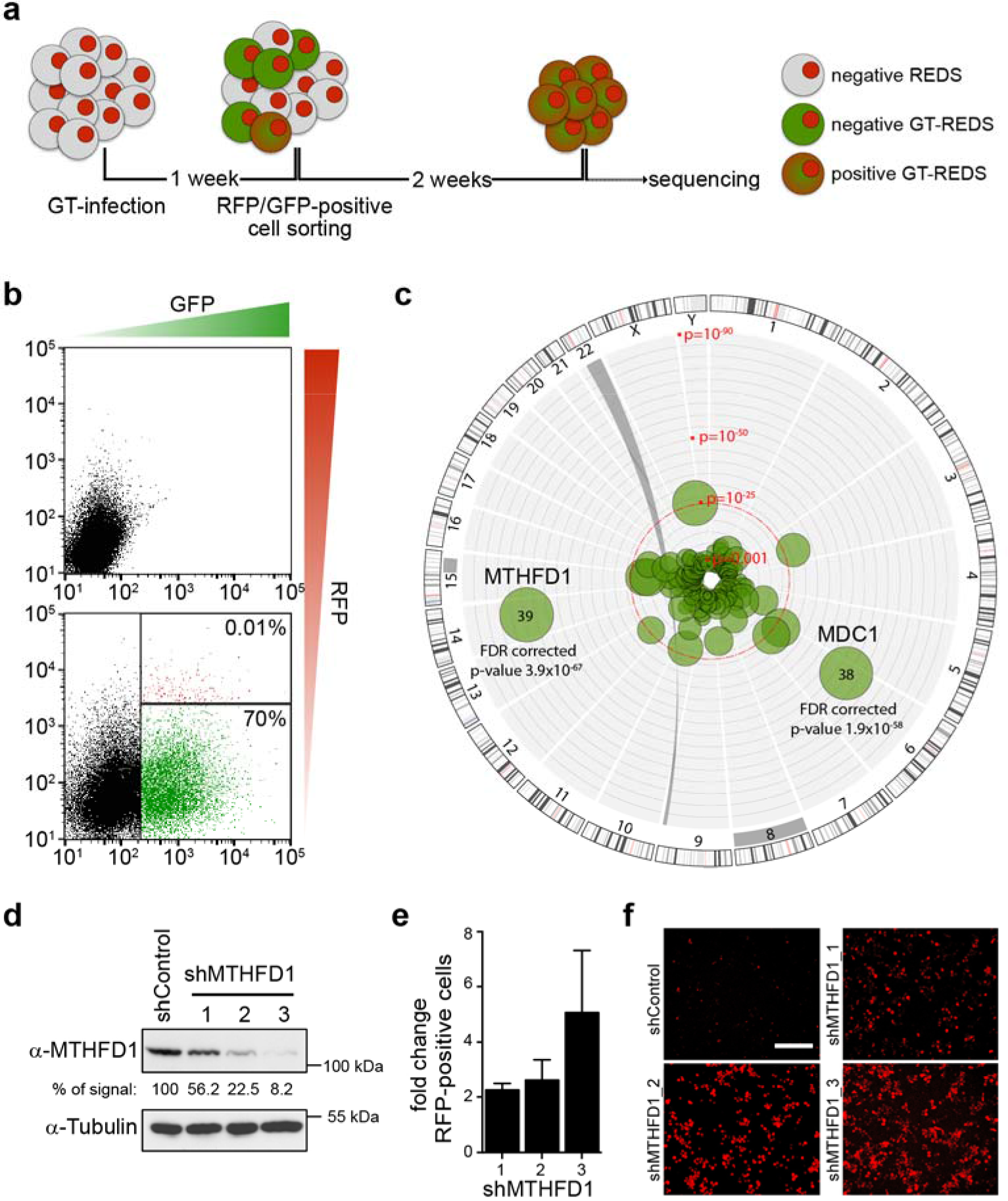
A genetic screen identifies MTHFD1 as a functional partner of BRD4. **a**, Schematic overview of the gene-trap based genetic screen. **b**, Representative panels of the applied FACS-sorting strategy. The upper panel shows non-infected REDS1 cells. The lower panel shows gene-trap infected REDS1 cells; three population can be distinguished: non-infected cells (black), infected GFP^+^ cells (green) and infected double positive (GFP^+^/RFP^+^) cells (red population: 0.01%). **c**, Circos-plot illustrating the hits from the gene-trap screen. Chromosome numbers are indicated on the outside, bubble size is proportional to the number of independent inactivating gene-trap integrations, distance from the center is proportional to significance (p values). **d**, Western blot showing MTHFD1 protein levels after downregulation with the indicated shRNAs in REDS1 cells. Numbers indicate the percentage of MTHFD1 protein remaining, tubulin was used as a loading control. **e**, Quantification of RFP^+^ cells from live-cell imaging of REDS1 cells treated with MTHFD1 shRNA. Three biological replicates were done for each experimental condition (mean±STD). **f**, Representative live-cell images of MTHFD1 knock down in REDS1 cells. Scale bar 100 μm.

In a second complementary screening approach, we used proteomics to identify BRD4 interactors in K-562, MOLM-13, MV4-11 and MEG-01 leukemia cell lines (Fig. 2a and Extended Data Fig. 3a). Only 13 proteins commonly interacted with BRD4 in all four cell lines. This set comprised several chromatin proteins like BRD3, LMNB1 and SMC3 and additionally included MTHFD1, the folate pathway enzyme identified in the genetic screen. The physical interaction between BRD4 and MTHFD1 was confirmed in the four leukemia cell lines (Fig. 2b) and all additional cell lines tested (Extended Data Fig. 3b-c). Recombinant MTHFD1 inhibited the interaction between BRD4 and acetylated histone peptides (Extended Data Fig. 3d), and the known acetylation site K56 in the MTHFD1 substrate pocket contributed to the binding (Extended Data Fig. 3e-h). In cellular pull-down assays all BRD4 isoforms interacted with full-length MTHFD1 but not with the dehydrogenase/cyclohydrolase or formyltetrahydrofolate-synthase domains alone (Extended Data Fig. 3i-k). The MTHFD1(K56A) mutation, which mimics the uncharged acetylated state, enhanced interaction with BRD4, while the mutation of the same residue to a changed arginine reduced the interaction. Consistently, the double bromodomain mutant GFP-BRD4 N140F/N433F showed drastically reduced binding to FLAG-MTHFD1 (Extended Data Fig. 3l). While BRD4 is localized almost exclusively to the nucleus, folate biosynthesis is considered to occur in the cytoplasm and mitochondria^20^. Only recently nuclear import of folate pathway enzymes including MTHFD1 has been described^21,22^. Nuclear versus cytosolic fractionation of HAP1, KBM7 and HEK293T cells indicated that a fraction of MTHFD1 can be detected in the nucleus in all three cell lines (Fig. 2c). We next tested in which cellular compartment the interaction between BRD4 and MTHFD1 occurred. MTHFD1 pull-downs in the cytosolic and nuclear fractions of HAP1, KBM7 and HEK293T revealed that the interaction with BRD4 was restricted to the nucleus (Fig. 2c). With the nucleus confirmed as the interaction site of BRD4 and MTHFD1, we asked whether the BRD4-MTHFD1 complex was chromatin-bound or rather found in the soluble nuclear faction. We prepared chromatin extracts comprising tightly DNA-bound proteins from HAP1 cells and checked for the presence of BRD4 and MTHFD1 by western blot. Both proteins were clearly detectable in the chromatin-bound fraction (Fig. 2d). To probe whether BRD4 recruits MTHFD1 to chromatin, we treated HAP1 cells with the small molecule degronimids dBET1^23^ and dBET6^4^. Two-hour treatment with these compounds resulted in the near-total ablation of BRD4 from chromatin. Under these conditions MTHFD1 was lost from chromatin, with remaining levels correlating with the amount of BRD4 (Fig. 2d), strongly suggesting that BRD4 is the sole factor recruiting MTHFD1 to chromatin. We further observed that the antifolate methotrexate (MTX) caused a similar depletion of chromatin-associated MTHFD1, while it did not affect BRD4 levels. A possible explanation is direct competition between BRD4 and MTX for binding to the MTHFD1 substrate pocked containing K56ac (Extended Data Fig. 3m). Importantly, BRD4 degradation or MTX treatment did not impair MTHFD1 nuclear localization (Fig. 2e), suggesting that while nuclear import of MTHFD1 is otherwise mediated, the interaction with BRD4 accounts for the recruitment of MTHFD1 to chromatin. To ensure cell-type independence, we confirmed that BRD4 degradation results in loss of MTHFD1 from chromatin in five additional cell lines (Fig. 2f-g).

**Figure 2.**
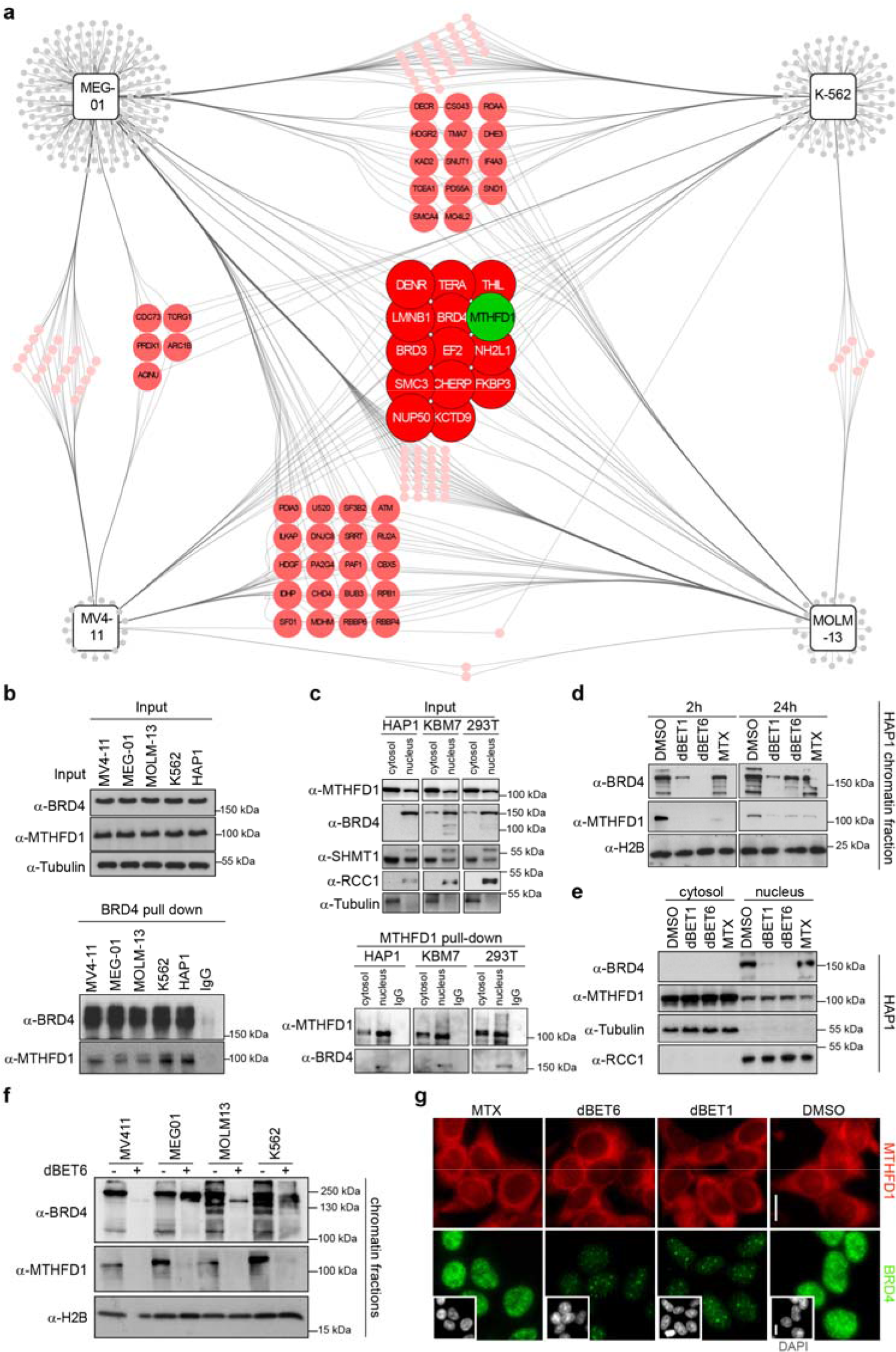
BRD4 recruits MTHFD1 to chromatin. **a**, BRD4 interactomes in MEG-01, K-562, MV4-11and MOLM-13 cell lines. Proteins are represented as circles, colors indicate the number of cell lines in which a particular interacting protein was detected. **b**, Western blot confirmation of the BRD4-MTHFD1 interaction in leukemia cell lines. **c**, Upper panel: Western blot following nuclear *vs* cytoplasmic fractionation in HAP1, KBM7 and HEK293T cell lines. RCC1 was used as nuclear loading control while tubulin was used as cytosolic loading control. Lower panel: Western blot following MTHFD1 pull-down in the different cell fractions. **d**, Western blot performed on chromatin-associated protein samples extracted from HAP1 cells treated with the indicated compounds for 2 h (dBET1: 0.5 μM; dBET6: 0.5 μM; MTX: 1 μM) or 24 h (dBET1: 0.5 μM; dBET6: 0.05 μM; MTX: 1 μM). H2B was used as loading control. **e**, Western blot for nuclear *vs* cytoplasmic protein levels in HAP1 cells treated for 24 h as above. **f**, Western blot from chromatin fractions of MEG-01, K-562, MV4-11 and MOLM-13 cells treated with dBET6 for 2 h. **g**, Immunofluorescence images of HeLa cells treated with the indicated compounds and stained for MTHFD1, BRD4, and DAPI (small inserts). Scale bar 10 μm.

Having characterized BRD4-dependent chromatin recruitment of MTHFD1, we mapped the genomic binding sites of the enzyme by ChIP-Seq experiments in HAP1 cells. MTHFD1 was bound to distinct genomic loci and binding was lost after 2 h treatment with dBET6 (Fig. 3a and Extended Data Fig. 4a-b). In line with the proteomic experiments, the vast majority of MTHFD1 binding sites overlapped with BRD4 binding sites at promoter and enhancers regions, where also H3K27ac was enriched (Fig. 3b-d and Extended Data Fig. 4c-e) suggesting a widespread role of MTHFD1 in transcriptional control. Transcriptome analysis of HAP1 cells revealed strong correlation of transcription changes following treatment with BRD4 inhibitors, degraders and antifolates, as well as between knock-down of BRD4 and of MTHFD1 (Fig. 3e and Extended Data Fig. 5a). Both MTHFD1 and BRD4 were enriched at promoters of genes that were downregulated following knock-down of either of these proteins (Fig. 3f and Extended Data Fig. 5b). The strong correlation between transcriptional effects of BET inhibitors and antifolates, as well as between knock-down of MTHFD1 and BRD4 is conserved in K-562 and A549 cells, suggesting cell type independence (Extended Data Fig. 6a-b).

**Figure 3.**
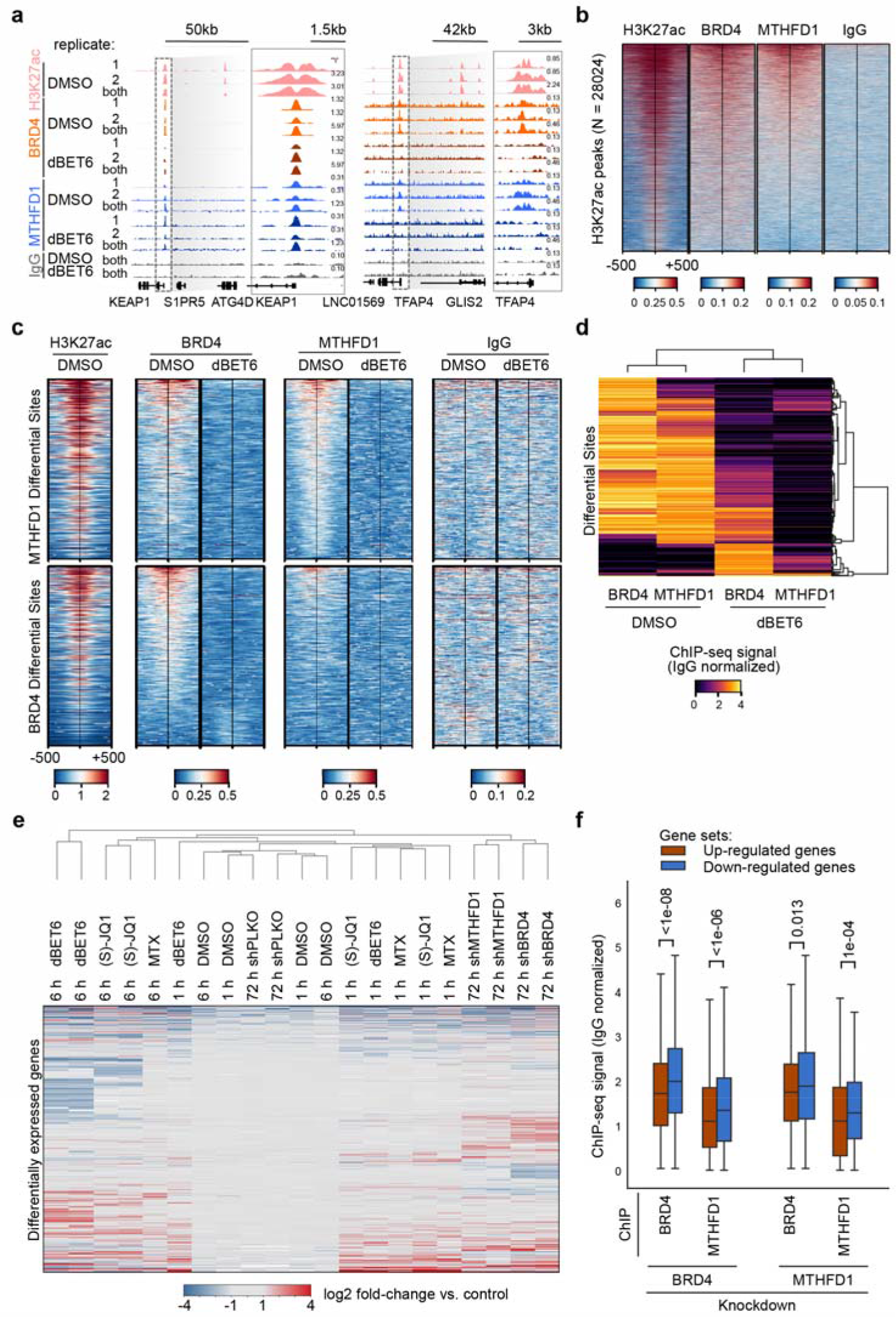
MTHFD1 regulates transcription by binding BRD4-occupied chromatin. **a**, Representative genome browser view of BRD4 and MTHFD1 binding in the H3K27ac-marked promoters of KEAP1 (left) and TFAP4 (right). All ChIP tracks were normalized to total mapped reads and the respective IgG control was subtracted from the merged replicate tracks. **b**, Enrichment of BRD4 and MTHFD1 ChIP signal in H3K27ac peaks. Peaks were sorted by H3K27ac abundance and data represent merged replicates in reads per basepair per million mapped reads. **c**, Enrichment of BRD4 and MTHFD1 in the top 500 differentially bound sites between dBET6 and DMSO treatment. **d**, Clustering of BRD4 and MTHFD1 abundance in the joint set of top 500 differentially bound sites between dBET6 or DMSO treatment. Hierarchical clustering with correlation as distance measurement was used. Values represent estimated factor abundance normalized by matched IgG signal. **e**, Heatmaps of transcriptome analysis of HAP1 cells treated with 0.1μM dBET6, 1μM (*S*)-JQ1, 1μM MTX, shRNAs targeting BRD4 or MTHFD1. Equal amount of DMSO, or non-targeting hairpins were used as respective control conditions. **f**, Integration of ChIP-Seq and RNA-Seq data in HAP1 cells. BRD4 and MTHFD1 binding at sites associated with genes which are up-(orange) or down-regulated (blue) upon knockdown of either BRD4 or MTHFD1. Values represent estimated factor abundance normalized by matched IgG signal and equality of distributions was assessed with the Kolmogorov-Smirnov test.

MTHFD1 is a C-1-tetrahydrofolate synthase that catalyzes three enzymatic reactions in folate metabolism, resulting in the interconversion of tetrahydrofolate, 10-formyltetrahydrofolate (10-CHO-THF), 5,10-methenyltetrahydrofolate (5,10-CH=THF) and 5,10-methylenetetrahydrofolate (5,10-CH2-THF). These folates are key intermediates of one carbon metabolism and provide activated C1 groups for the biosynthesis of purines, pyrimidines and methionine. Biosynthesis of these three major classes of C1 metabolism products is considered to occur predominantly in the cytoplasm and mitochondria of mammalian cells^20^, but we detected the majority of enzymes required for nucleotide biosynthesis also in the tightly chromatin-associated protein fraction in K-562 and HAP1 cells (Extended Data Fig. 7a and Extended Data Table 2). These data indicate that nucleotide biosynthesis may also occur in a chromatin environment, and we asked whether inhibition of BRD4 or MTHFD1 altered nuclear metabolite composition. We knocked-down either BRD4 or MTHFD1, isolated nuclei and analyzed their composition in a metabolomics approach relative to a non-targeting control hairpin. In total, we detected 2851 metabolites, of which over 400 were significantly changed in one of the conditions. We observed strong correlation between the nuclear metabolomes in BRD4 and MTHFD1 knock-down conditions, particularly for metabolites in the purine, pyrimidine and methionine biosynthesis pathways (Extended Data Fig. 7b-d). The direct MTHFD1 product 5,10-CH2-THF and most of the precursors for *de novo* nucleoside and nucleotide biosynthesis were depleted in both conditions, whereas the vast majority of free bases, nucleosides and nucleotides were increased. Furthermore, the trend for similar changes in nuclear metabolite composition was also noticeable in cells treated with small molecules dBET1 and MTX (Extended Data Fig. 7e). BET inhibitors and MTX caused highly correlated characteristic changes specifically in the nuclear folate pool that were not observed with other cytotoxic compounds (Extended Data Fig. 7f). Overall, a common nuclear metabolite signature for the inhibition of MTHFD1 and of BRD4 is evident, indicating a crosstalk between BRD4-dependent epigenetic regulation and folate metabolism.

Based on the similarities in nuclear metabolite composition following loss of MTHFD1 and BRD4, we speculated that antifolates might synergize with BRD4 inhibitors. To test this hypothesis, we treated REDS cells with (*S*)-JQ1 and MTX alone and in combination. Co-treatment with MTX remarkably amplified the basal RFP signal given by low doses of (*S*)-JQ1 alone (Fig. 4a). These results indicate that the chromatin remodeling process can be enhanced when inhibiting BRD4 and MTHFD1 together, emphasizing the fundamental role of folate metabolites in epigenetic regulation. We next tested whether the synergism also impacts cancer cell survival and selected six cell lines including four cell lines described as not sensitive to BRD4 inhibition, plus KBM7 and HAP1 cells (Extended Data Fig. 8a). Dose response curves confirmed the low sensitivity of these cell lines to (*S*)-JQ1 treatment and a moderate to low sensitivity to MTX treatment (Extended Data Fig. 8b). In contrast to the poor response to (*S*)-JQ1 and MTX individual treatments, the combination of both drugs efficiently impaired cell viability in all six cell lines tested (Fig. 4b and Extended Data Fig. 8c). Toxicity was observed at concentrations without any single-agent activity, indicating strong synergism between the two treatments, also confirmed by calculating synergy indices according to the Bliss independence model ^24^ (Extended Data Fig. 8d). To exclude possible off-target effects of MTX, we treated the cell line showing the strongest drug synergism, A549, with shRNA for MTHFD1 and demonstrated increased sensitivity to (*S*)-JQ1 (Extended Data Fig. 8e). We then proved that BET bromodomain inhibitors can be combined with antifolates *in vivo* to specifically inhibit cancer cell proliferation without exerting general toxicity. When we treated an A549 xenograft mouse model^25^ with MTX and (*S*)-JQ1 alone and in combination, tumor growth was not impaired by either of the individual compounds, but arrested when the two inhibitors were given together (Fig. 4d-f).

**Figure 4.**
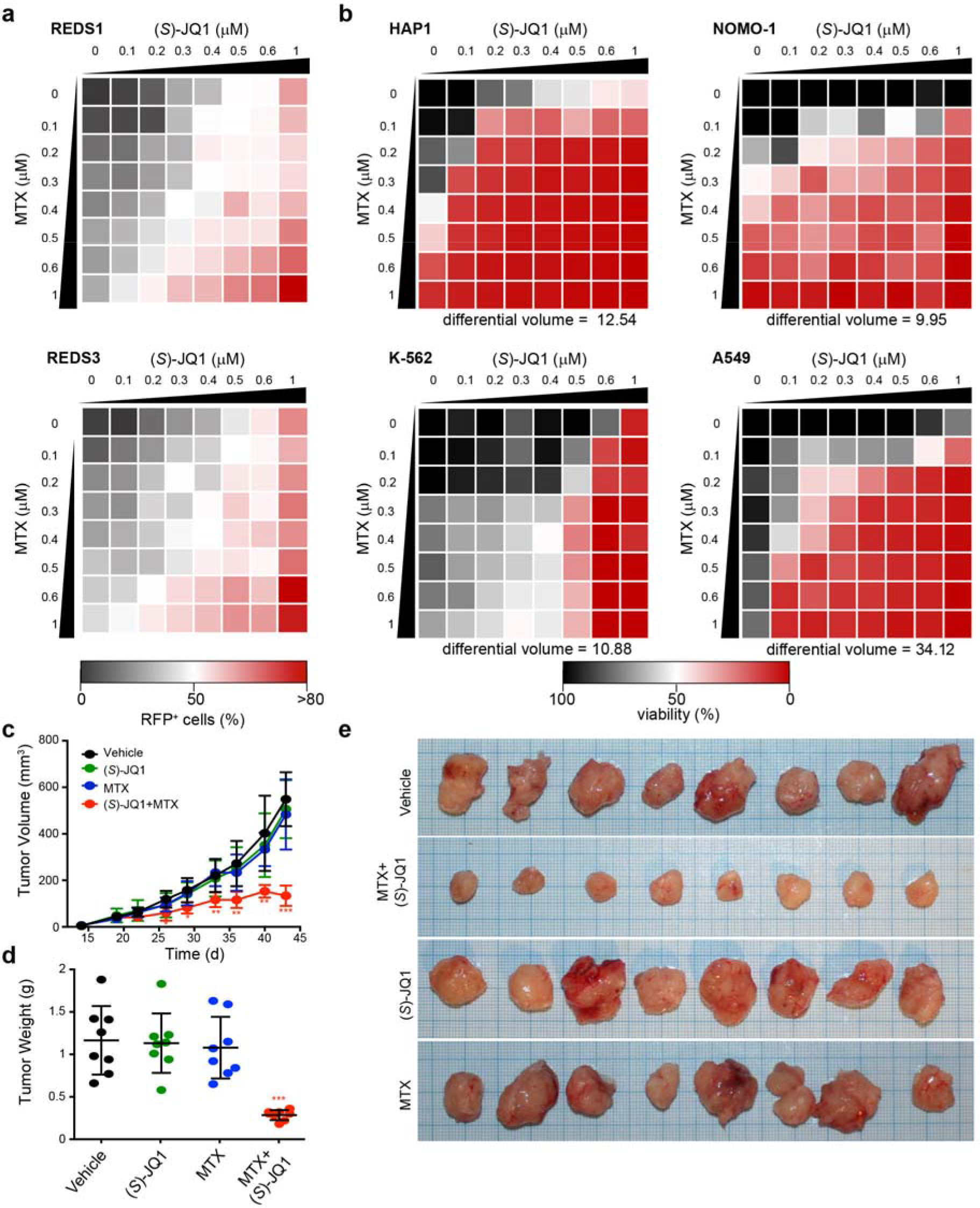
BET bromodomain inhibitors synergize with antifolates to impair cancer cell growth. **a**, Dose response matrix displaying REDS1 and REDS3 RFP-positive cells treated with the indicated concentrations of (S)-JQ1 and MTX alone or in combination. Means from two biological replicates. **b**, Dose response matrices displaying cell viability of HAP1, NOMO-1, K-562 and A549 treated for 72 h with (*S*)-JQ1 and MTX alone or in combination. Means from two biological replicates. Differential volume indicates the sum of all deviations from Bliss additivity over the dose response matrix. **c**, Tumor volumes from a A549 xenograft mouse model treated five times per week with 30 mg/kg (*S*)-JQ1 and/or twice weekly with 25 mg/kg MTX from day 19. Means and standard deviations from eight mice per group. Asterisks indicate significance (* p<0.05; ** p<0.005; *** p<0.0001) **d**, Weight and **e**, images of tumors at the end of the experiment (day 43).

In contrast to nuclear ATP and acetyl-CoA biosynthesis^26,27^, the role of folate pathway enzymes in the control of gene expression had not been studied. Here we discovered and characterized a novel transcriptional role for nuclear folate metabolism. We show that at distinct genomic loci control of gene expression by BRD4 relies on the physical interaction with MTHFD1 and both proteins are required to establish and maintain a pool of folate metabolites in the nucleus. Interestingly, we identified not only MTHFD1 but also the other enzymes of *de novo* nucleotide biosynthesis pathway physically bound to chromatin. Nucleotides are also the metabolites we find most dramatically changed following loss of either BRD4 or MTHFD1. Nucleotide stress is known to impair P-TEFb activation and transcriptional elongation via upregulation of HEXIM1^28^. Therefore, we hypothesize that BRD4-dependent MTHFD1 recruitment is needed to support local nucleotide availability for full transcriptional activation of certain BRD4-target genes needed for cancer cell proliferation.

Due to its fundamental role in cell proliferation *via* nucleic acid biosynthesis, folate metabolism has been widely investigated in cancer biology. Several small molecules, including MTX, that target different enzymes of the folate pathway^29^ have been developed. These antifolates are considered to act by inhibiting cell division, DNA/RNA synthesis and repair and protein synthesis^30^. Our findings suggest an additional ‘targeted’ mode of these classical chemotherapeutic drugs, which should enable better stratification and treatment regimens for cancer patients. Furthermore, we suggest combination of antifolates with BRD4 inhibitors as possible treatment for particularly aggressive cancers, opening the avenue for more successful therapies.

## METHODS

### Cell culture and transfection

KBM7 (human chronic myelogenous leukemia), MV4-11 (biphenotypic B myelomonocytic leukemia), MEG-01 (human chronic myelogenous leukemia), K-562 (human chronic myelogenous leukemia) and HAP1 (KBM7-derived) cell lines were cultured in Iscove’s Modified Dulbecco’s Medium (IMDM, Gibco), supplemented with 10% Fetal Bovine Serum (FBS; Gibco). HEK293T (human embryonic kidney) and HeLa (cervix adenocarcinoma) cell lines were cultured in Dulbecco’s Modified Eagles Medium (DMEM, Gibco) supplemented with 10% FBS. MOLM-13 (human acute monocytic leukemia), NOMO-1 (human acute monocytic leukemia) and A549 (lung carcinoma) cell lines were cultured in RPMI-1640 (Roswell Park Memorial Institute, Gibco) supplemented with 10% FBS. All the mentioned cell lines were incubated in 5% CO_2_ atmosphere at 37°C.

HEK293T cells were transfected with Lipofectamine 2000 (Invitrogen) according to the manufacturer’s instructions.

The Retroviral Gene-Trap vector (pGT-GFP; see below) was a kind gift from Dr. Sebastian Nijman, Group Leader at the Cell biology, Signaling, Therapeutics Program, Ludwig Cancer Research (Oxford, UK). GFP-MTHFD1 plasmid was a kind gift from Prof. Patrick Stover, Director of the Division of Nutritional Sciences, Cornell University (Ithaca, NY).

### Western blot and immunoprecipitation

For Western blot, proteins were separated on polyacrylamide gels with SDS running buffer (50 mM Tris, 380 mM Glycine, 7 mM SDS) and transferred to nitrocellulose blotting membranes. All membranes were blocked with blocking buffer (5% (m/v) milk powder (BioRad) in TBST (Tris-buffered saline with Tween: 50 mM Tris (tris (hydroxymethyl)aminomethane), 150 mM NaCl, 0,05% (v/v) Tween 20, adjusted to pH 7.6). Proteins were probed with antibodies against BRD4 (ab128874, 1:1000, Abcam), Actin (ab16039, 1:1000, Abcam), MTHFD1 (ab70203, Abcam; H120, Santa Cruz; A8, Santa Cruz. All used at 1:1000), GFP (G10362, 1:1000, Life Tecnhology), RCC1 (C-20, 1:1000, Santa Cruz), β-Tubulin (T-4026, 1:1000, Sigma), SHMT1 (ab186130, 1:1000, Abcam) and H2B (ab156197, 1:1000, Abcam) and detected by HRP (horseradish peroxidase) conjugated donkey anti-rabbit IgG antibody (ab16284, 1:5000, Abcam) or donkey anti-mouse IgG antibody (Pierce) and visualized with the Pierce ECL Western Blotting substrate (Amersham), according to the provided protocol.

For immunoprecipitation, 1 mg of protein extract was incubated overnight at 4°C with 10 μl Protein A or G beads (Life Technologies) preincubated for 2 hours at 4°C with 1 μg of BRD4 (ab128874, Abcam), MTHFD1 (A8, Santa Cruz) or GFP (G10362, Life Technology) antibodies.

### Immunofluorescence, metaphase spread and live cell imaging

For immunofluorescence, cells were grown on coverslips precoated with Polylysine (Sigma). After the desired treatment, cells were washed with PBS and fixed with cold MeOH for at least 24 hours. Blocking was performed in PBS/3% bovine serum albumin (BSA)/0.1% Triton for 30 minutes. Coverslips were incubated first with primary antibodies for 30 minutes at room temperature (MTHFD1 H120, Santa Cruz; BRD4 ab128874, Abcam) and then with secondary antibodies (Alexa Fluor 488 Goat Anti-Rabbit and Alexa Fluor 546 Donkey Anti-Mouse, Thermo Fisher Scientific) for 30 minutes in the dark. Finally, they were washed and incubated with DAPI (4,6-diamidino-2-phenylindole) for 5 minutes at room temperature in the dark. Three PBS washing steps were done to remove the excess of antibodies and DAPI, and coverslips were mounted with Propyl-Gallate (Sigma) on slides. Pictures were taken with a Leica DMI6000B inverted microscope and 63X oil objective and analyzed with Fiji (ImageJ).

Metaphase spreads were obtained from cells treated overnight with nocodazole (250 ng/ml). Cells were washed with PBS and the pellet was resuspended carefully in prewarmed (37 °C) 0.56% KCl and incubated at 37 °C for 10 minutes. Cells were centrifuged at 800g for 5 minutes and resuspended in ice-cold and freshly prepared fixative (MeOH:Acetic Acid/3:1). Metaphase spreads were prepared dropping 30 μl of fixed cell suspension on glass slides previously dipped in ice-cold 40% MeOH and dried. The fixation solution was rapidly evaporated placing the slides at 75 °C for 10 seconds. Finally, chromatin was stained with DAPI for 5 minutes; three PBS washing steps were done to remove the excess of DAPI, and coverslips were mounted with Propyl-Gallate (Sigma) on slides. Pictures were taken with a Leica DMI6000B inverted microscope and 63X oil objective and analyzed with Fiji (ImageJ).

Live cell imaging pictures were taken from cells seeded on clear flat bottom 96-well or 384-well plates (Corning), with the Operetta High Content Screening System (PerkinElmer), 20X objective and non-confocal mode. RFP quantification was done using the basic PerkinElmer software for nuclei detection and analysis, adapted for the nucleus diameter range of the specific cell line used (KBM7, 13 μm). Only RFP positive nuclei were detected and counted.

### Cell cycle assay

For cell cycle analysis, 1 million cells were fixed with 70% ethanol for 24 hours, washed with PBS/1% BSA/0.1% Tween and incubated with RNase for 30 minutes. Nuclei were stained with 5 μg/ml PI (propidium iodide, Sigma) for 10 minutes prior to FACS analysis (BD FACSCalibur Flow Cytometer).

### RNA extraction and RT-PCR

RNA extraction was performed with TRIzol Reagent (Life Technologies) according to the standard protocol and Reverse Transcription (RT) was performed using the High Capacity cDNA Reverse Transcription Kit (Applied Biosystems).

QPCR was performed using the Power SYBR Green Master mix (Invitrogen) as described in the manufacture’s protocol.

QPCR primers used:

Actin (Sigma; forward 5’-ATGATGATATCGCCGCGCTC, reverse 5’-CCACCATCACGCCCTGG).
BRD1 (Sigma; forward 5’-GAAGAAGCAGTTTGTGGAGC, reverse 5’-GCAGTCTCAGCGAAGCTCAC).
BRD2 (Sigma; forward 5’-GCTTGGGAAGACTTTGTTGG, reverse 5’-TGTCAGTCACCAGGCAGAAG)
BRD3 (Sigma; forward 5’-AAGAAGAAGGACAAGGAGAAGG, reverse 5’-CTTCTTGGCAGGAGCCTTCT).
BRD4 (Sigma; forward 5’-CAGGAGGGTTGTACTTATAGCA, reverse 5’-CTACTGTGACATCATCAAGCAC).
BRDT (Sigma; forward 5’-TCAAAGATCCCGATTGAACC, reverse 5’-CGGAAAGGTACTTGGGACAA)

Real-time amplification results were normalized to the endogenous housekeeping gene Actin. The relative quantities were calculated using the comparative CT (Cycle Threshold) Method (ΔΔCT Method).

### Gene-Trap genetic screening

pGT-GFP contains an inactivated 3’ LTR, a strong adenoviral (Ad40) splice-acceptor site, the GFP coding sequence and the SV40 polyadenylation signal. The Gene-trap virus was produced by transfection of 293T cells in T150 dishes with pGT-GFP combined with retroviral packaging plasmids. The virus-containing supernatant was collected after 30, 48 and 72 hours of transfection and concentrated using ultracentrifugation for 1.5 hours at 24100 rpm in a Beckman Coulter Optima L-100 XP ultracentrifuge using an SW 32 Ti rotor.

For each replicate, 20 million REDS1 cells were mutagenized in a 24-well plate, seeding 1 million cells per well and using spin-infection for 45 minutes at 2000 rpm. GT-infected cells were assessed by FACS to determine the percentage of infection (percentage of GFP positive cells). If such percentage was above 70%, REDS1 GFP/RFP double positive cells were sorted and left in culture for 2 weeks to get the sufficient number of cells to process for the DNA library preparation.

### Gene-Trap analysis

Raw sequencing data were aligned to human reference genome hg19 (UCSC hg19 build) using bowtie2 (version 2.2.4) with default parameter. Reads that did not meet the following criteria were removed: (1) have a reported alignment - “mapped reads”; (2) have a unique alignment; (3) have a mapping quality (MAPQ) higher than 20. Duplicate reads were marked and discarded with Picard (version 1.111). Insertions in close proximity (1 or 2 base pairs distance from each other) were removed to avoid inclusion of insertions due to mapping errors. Insertions were annotated with gene build GRCh37.p13 (ENSEMBL 75 - release February 2014) using bedtools (version 2.10.1) and custom scripts. The canonical transcripts (according to ENSEMBL) for each gene were used as a reference gene model to count insertions falling with exons, introns or intragenic. Insertions were considered mutagenic or disruptive to the gene if they occurred within exons irrespective of their orientation to the corresponding gene or if they were located within introns in sense orientation ^31^. Insertions in antisense direction in respect to the gene orientation were considered silent. All mutagenic insertions were summarized independently for each gene. For each gene a one-sided Fisher’s exact-test was applied to estimate a significant enrichment of insertions over an unselected control data set. Resulting p-values were adjusted for false discovery rate (FDR) using Benjamini-Hochberg procedure. A cutoff of 5% FDR was considered significantly enriched. Insertion plots were drawn with R statistics software and circos plots were produced using circos plots ^32^.

### Cell sorting

RFP/GFP double positive cell sorting was performed using the FACSAria (BD Biosciences) sorter. Gates for positive or negative RFP or GFP populations were done using the appropriate positive or negative controls. RFP/GFP double positive cells were sorted 7 days after GT infection.

### DNA library preparation

DNA was extracted from 30 million GFP/RFP double positive REDS1 cells using the Genomic DNA isolation QIAamp DNA mini kit (Qiagen). 4 μg of DNA were digested with NlaIII or MseI (4 digestions each enzyme). After spin column purification (Qiagen), 1 μg of digested DNA was ligated using T4 DNA ligase (NEB) in a volume of 300 μl (total of 4 ligations). The reaction mix was purified and retroviral insertion sites were identified via an inverse PCR protocol adapted to next generation sequencing ^33^.

### FISH assay

The RFP specific probe (RFP_probe) was PCR performed using RFP specific primers (Sigma; forward 5’-CGGTTAAAGGTGCCGTCTCG, reverse 5’-AGGCTTCCCAGGTCACGATG) and labeled using dig-dUTP (DIG Nick Translation Mix, Roche). The FISH assay procedure was performed as previously described ^17^.

### AlphaLISA assay

We performed the Amplified Luminescent Proximity Homogenous Assay (AlphaLISA^©^), a homogenous and chemiluminescence-based method, to explore the direct interaction of BRD4 and acetylated substrates.

Briefly, in this assay, the biotinylated MTHFD1 acetylated peptides (possible substrates) are captured by streptavidin-coupled donor beads. GST-tagged BRD4 (produced as previously described ^17^) is recognized and bound by an anti-GST antibody conjugated with an acceptor bead. In case of interaction between BRD4 and one acetylated peptide, the proximity between the partners (< 200 nm) allows that the excitation (680 nm wavelength) of a donor bead induces the release of a singlet oxygen molecule (^1^O_2_) that then triggers a cascade of energy transfer in the acceptor bead, resulting in a sharp peak of light emission at 615 nm.

GST-BRD4 and each of the MTHFD1 acetylated peptides were incubated together. After 30 minutes, GSH (Glutathione) Acceptor beads (PerkinElmer) were added and after another incubation time of 30 minutes, Streptavidin-conjugated donor beads (PerkinElmer) were added. The signal (alpha counts) was read by the EnVision 2104 Multilabel Reader (PerkinElmer).

### Preparation of nuclear cell extracts for proteomics

Nuclear extract was produced from fresh cells grown at 5 millions of cells/mL. Cells were collected by centrifugation, washed with PBS and resuspended in hypotonic buffer A (10 mM Tris-Cl, pH 7.4, 1.5 mM MgCl2, 10 mM KCl, 25 mM NaF, 1 mM Na_3_VO_4_, 1 mM DTT, and protease inhibitor cocktail (cOmplete, Roche)). After ca. 3 minutes, cells were spun down and resuspended in buffer A and homogenized using a Dounce homogenizer. Nuclei were collected by centrifugation in a microfuge for 10 minutes at 3300 rpm, washed with buffer A and homogenized in one volume of extraction buffer B (50 mM Tris-Cl, pH 7.4, 1.5 mM MgCl_2_, 20 % glycerol, 420 mM NaCl, 25 mM NaF, 1 mM Na_3_VO_4_, 1 mM DTT, 400 Units/ml DNase I, and protease inhibitor cocktail). Extraction was allowed to proceed under agitation for 30 minutes at 4 °C before the extract was clarified by centrifugation at 13000g. The extract was diluted 3:1 in buffer D (50 mM Tris-Cl, pH 7.4 (RT), 1.5 mM MgCl_2_, 25 mM NaF, 1 mM Na_3_VO_4_, 0.6% NP40, 1 mM DTT, and protease inhibitor cocktail), centrifuged again, and aliquots were snap frozen in liquid nitrogen and stored at −80°C.

### Immunopurification (IP-MS) and nanoLC-MS analysis

Anti-BRD4 (A301-985A, Bethyl Labs) antibody (50 μg) was coupled to 100 μl AminoLink resin (Thermo Fisher Scientific). Cell lysate samples (5 mg) were incubated with prewashed immuno-resin on a shaker for 2 hours at 4 °C. Beads were washed in lysis buffer containing 0.4% Igepal-CA630 and lysis buffer without detergent followed by two washing steps with 150 mM NaCl. Samples were processed by on-bead digest with Lys-C and Glycine protease before they were reduced, alkylated and digested with Trypsin.

The nano HPLC system used was an UltiMate 3000 HPLC RSLC nano system (Thermo Fisher Scientific, Amsterdam, Netherlands) coupled to a Q Exactive mass spectrometer (Thermo Fisher Scientific, Bremen, Germany), equipped with a Proxeon nanospray source (Thermo Fisher Scientific, Odense, Denmark).

The Q Exactive mass spectrometer was operated in data-dependent mode, using a full scan (m/z range 350-1650, nominal resolution of 70 000, target value 1E6) followed by MS/MS scans of the 12 most abundant ions. MS/MS spectra were acquired using normalized collision energy 30%, isolation width of 2 and the target value was set to 5E4. Precursor ions selected for fragmentation (charge state 2 and higher) were put on a dynamic exclusion list for 30 s. Additionally, the underfill ratio was set to 20% resulting in an intensity threshold of 2E4. The peptide match feature and the exclude isotopes feature were enabled.

### MS data analysis (proteomics)

For peptide identification, the RAW-files were loaded into Proteome Discoverer (version 1.4.0.288, Thermo Scientific). All hereby created MS/MS spectra were searched using Mascot 2.2.07 (Matrix Science, London, UK) against the human swissprot protein sequence database. The following search parameters were used: Beta-methylthiolation on cysteine was set as a fixed modification, oxidation on methionine. Monoisotopic masses were searched within unrestricted protein masses for tryptic peptides. The peptide mass tolerance was set to ±5 ppm and the fragment mass tolerance to ±30 mmu. The maximal number of missed cleavages was set to 2. For calculation of protein areas Event Detector node and Precursor Ions Area Detector node, both integrated in Thermo Proteome Discoverer, were used. The result was filtered to 1% FDR using Percolator algorithm integrated in Thermo Proteome Discoverer. Additional data processing of the triplicate runs including label-free quantification was performed in MaxQuant using the Andromeda search engine applying the same search parameters as for Mascot database search. For subsequent statistical analysis Perseus software platform was used to create volcano plots, heat maps and hierarchical clustering.

### Molecular modeling

For calculating the binding affinity of MTHFD1(K56ac) towards BRD4, six crystal structures of BRD4 co-crystallized with any peptide were downloaded from the RCSB Protein Databank (PDB; www.rcsb.org) ^34^. The X-ray structures were prepared using the QuickPrep protocol of the MOE software package. With that, hydrogens and missing atoms were added, charges were calculated, protonation states optimized and clashes and strain were removed by performing a short energy minimization. Prior to mutating the co-crystallized peptide into the MTHFD1(K56ac), the crucial interaction of the acetylated Lys with Asn140 was restrained. The virtual mutations as well as the affinity and stability calculations were performed using the Protein Design tools (Residue Scan with default settings) of the MOE software package.

For predicting the binding of Methotrexate (MTX) to MTHFD1 (acetylated and unacetylated), the X-ray structure 1A4I was prepared with the QuickPrep protocol of MOE. As the binding pocket of 1A4I is highly solvated, water molecules might interfere with MTX binding during the docking run. Therefore, water molecules were removed for all calculations. For the comparison of binding acetylated *vs* unacetylated MTHFD1, the prepared crystal structure was acetylated using the Protein Builder in MOE, followed by a short energy minimization of the mutated residue. Furthermore, MTX was prepared and protonated in MOE. A conformational analysis using the LowModeMD method with default settings provided 37 different MTX conformations. These 37 conformations were docked into the acetylated and unacetylated structures of MTHFD1 using the induced fit docking protocol in MOE with default settings.

Interaction fingerprints of the docked structures were calculated using the PLIF tool in MOE.

### Chromatin purification and LC-MS/MS analysis

Cell fractionation and chromatin enrichment was carried out as previously described ^35^ with some adaptations. Briefly, for 100 million cells, the chromatin enriched pellet was taken up in 250 μl benzonase digestion buffer (15mM HEPES, 1mM EDTA, 1mM EGTA, 0.1% NP40, protease inhibitor cocktail (cOmplete, Roche)) after washing, and sonicated for 120 seconds on the Covaris S220 focused-ultrasonicator with the following settings: Peak Power 140; Duty-Factor 10.0; Cycles/Burst 200. After addition of 0.25U benzonase and 2.5 μg RNase, the chromatin was incubated for 40 minutes at 4°C on a rotary shaker. 2x SDS lysis buffer (100mM HEPES, 4% SDS, 2 mM PMSF and protease inhibitor cocktail (cOmplete, Roche)) was added to the samples in a 1:1 ratio and incubated for 10 minutes at room temperature followed by 5 minutes denaturation at 99 °C. After centrifugation at 16.000 g for 10 minutes at room temperature, the supernatant was transferred to a new tube. MS sample preparation was perform using the FASP protocol as previously described ^36,37^. Reverse-phase chromatography at high and low pH was performed for two-dimensional peptide separation prior to MSMS analysis. Peptides were purified using solid-phase extraction (SPE) (MacroSpin Columns, 30-300μg capacity, Nest Group Inc. Southboro, MA, USA) and reconstituted in 23 μL 5% acetonitrile, 10 mM ammonium formate. An Agilent 1200 HPLC system (Agilent Biotechnologies, Palo Alto, CA) equipped with a Gemini-NX C18 (150 × 2 mm, 3 μm, 110 Å, Phenomenex, Torrance, US) column was used for the first dimension of liquid chromatography. Peptides were separated into 20 time based fractions during a 30 min gradient ranging from 5 to 90% acetonitrile containing 10 mM ammonium formate, pH 10, at a flow rate of 100 μL/min. Samples were acidified by the addition of 5 μL 5% formic acid. Solvent was removed in a vacuum concentrator, and samples were reconstituted in 5% formic acid.

Mass spectrometric analyses were performed on a Q Exactive mass spectrometer (ThermoFisher, Bremen, Germany) coupled online to an Agilent 1200 series dual pump HPLC system (Agilent Biotechnologies, Palo Alto, CA). Samples were transferred from the thermostatted autosampler (4 °C) to a trap column (Zorbax 300SB-C18 5 μm, 5 × 0.3 mm, Agilent Biotechnologies, Palo Alto, CA, USA) at a constant flow rate of 45 μL/min. Analyte separation occurred on a 20 cm 75 μm inner diameter analytical column, that was packed with Reprosil C18 (Dr. Maisch, Ammerbuch-Entringen, Germany) in house. The 60-minute gradient ranged from 3 % to 40 % organic phase at a constant flow rate of 250 nL/min. The mobile phases used for the HPLC were 0.4 % formic acid and 90 % acetonitrile plus 0.4 % formic acid for aqueous and organic phase, respectively. The Q Exactive mass spectrometer was operated in data-dependent mode with up to 10 MSMS scans following each full scan. Previously fragmented ions were dynamically excluded from repeated fragmentation for 20 seconds. 100 ms and 120 ms were allowed as the maximum ion injection time for MS and MSMS scans, respectively. The analyzer resolution was set to 70,000 for MS scans and 35,000 for MSMS scans. The automatic gain control was set to 3 × 106 and 2 × 105 for MS and MSMS, respectively, to prevent the overfilling of the C-trap. The underfill ratio for MSMS was set to 6 %, which corresponds to an intensity threshold of 1 × 105 to accept a peptide for fragmentation. Higher collision energy induced dissociation (HCD) at a normalized collision energy (NCE) of 34 was employed for peptide fragmentation and reporter ion generation. The ubiquitous contaminating siloxane ion Si(CH_3_)_2_O)_6_ was used as a single lock mass at m/z 445.120024 for internal mass calibration.

### MS data analysis (chromatin fraction)

The acquired raw MS data files were processed as previously described ^37^. The resultant peak lists were searched against the human SwissProt database version 20150601 with the search engines Mascot (v.2.3.02) and Phenyx (v.2.5.14).

For TMT quantitation the isobar R package was used ^38^. As the first step of the quantitation, the reporter ion intensities were normalized in silico to result in equal median intensity in each TMT reporter channel. Isobar statistical model considers two P-values: *P-value sample* that compares the abundance changes due to the treatment to the abundance changes seen between biological replicates and *P-value ratio* that models for noise/variability in mass spectrometry data collection. *P-value ratio* was further corrected for false discovery rate (FDR). Abundance of a protein was considered to be changed significantly if both *P-value sample* and FDR corrected *P-value ratio* were less than 0.05.

### Preparation of nuclear cell extracts for metabolomics

Nuclei were extracted by hypotonic lysis. Briefly, intact cells treated (as indicated in the results section) were washed twice with cold PBS and incubated on ice for 10 minutes with hypotonic lysis buffer (10 mM HEPES, pH 7.9, with 1.5 mM MgCl_2_, 10 mM KCl and protease inhibitor cocktail (cOmplete, Roche); buffer-cells volume ratio 5:1). Pellet was gently resuspended three times during the incubation. Nuclei were collected by centrifugation (420 g X 5 minutes) and immediately snap frozen.

The metabolomic assay and data analysis was performed by Metabolomic Discoveries (http://www.metabolomicdiscoveries.com; Germany). Briefly, LC-QTOF/MS-based non-targeted metabolite profiling was used to analyses nuclear metabolites in the range of 50-1700 Da, with an accuracy up to 1-2 ppm and a resolution of mass/Δmass=40.000. Metabolites measured in the LC are annotated according to their accurate mass and subsequent sum formula prediction. Metabolites that were not annotated in the LC-MS-analyses are listed according to their accurate mass and retention time.

### Metabolite set enrichment analysis

Metabolite set enrichment analysis (MSEA) ^39^ was performed using the online tool MetaboAnalyst ^40^ (http://www.metaboanalyst.ca/). Briefly, for each pre-defined functional group a fold-change is computed between the observed number of significantly altered metabolites (considering both up-and down-regulation, t-test with p-value < 0.05) and random expectation, as well as a corresponding pvalue (using Fisher’s exact test). In Figure S7D, we report all functional groups that contain at least one altered metabolite. Compounds with matching KeGG IDs were used in the comparison.

### Folate extraction and LC MS/MS analysis

In order to quantify folates in the nuclear and cytosolic fractions, 20 millions of HAP1 cells per condition were washed twice with cold PBS, and collected into 50 ml falcon tube by centrifugation for 5 minutes at 280 g and 4 °C. Cell lysis was performed on ice in the dark by incubating cell pellets with 1:5 hypotonic lysis buffer for 10 minutes. Nuclei were collected by centrifugation for 5 minutes at 420 g and 4 °C. Supernatants (cytosolic fractions) were also collected. Both fractions were immediately snap frozen.

For nucleus samples, 10 μL of ISTD mixture was added to nucleus pellet in 1.5 mL Eppendorf tube followed by addition of 145 μL of ice-cold extraction solvent (10 mg/mL ascorbic acid solution in 80% methanol, 20% water, v/v). The samples were vortexed for 10 seconds, afterwards incubated on ice for 3 min and vortexed again for 10 seconds. After centrifugation (14000 rpm, 10 min, 4 °C), the supernatant was collected into HPLC vials. The extraction step was repeated and combined supernatants were used for LC-MS/MS analysis.

For cytoplasm samples, 10 μL of ISTD mixture was added to 75 μL of cytoplasm 1.5 mL Eppendorf tube followed by addition of 215 μL of ice-cold extraction solvent (10 mg/mL ascorbic acid solution in 80% methanol, 20% water, v/v). The samples were vortexed for 10 seconds, afterwards incubated on ice for 3 min and vortexed again for 10 seconds. After centrifugation (14000 rpm, 10 min, 4 °C), the supernatant was collected into HPLC vials and used for LC-MS/MS analysis.

An Acquity UHPLC system (Waters) coupled with Xevo TQ-MS (Waters) triple quadrupole mass spectrometer was used for quantitative analysis of metabolites. The separation was conducted on an ACQUITY HSS T3, 1.8 μm, 2.1×100 mm column (Waters) equipped with an Acquity HSS T3 1.8 μM Vanguard guard column (Waters) at 40 °C. The separation was carried out using 0.1% formic acid (v/v) in water as a mobile phase A, and 0.1% formic acid (v/v) in methanol as a mobile phase B. The gradient elution with a flow rate 0.5 mL/min was performed with a total analysis time of 10 min. The autosampler temperature was set to 4 °C. For detection, Waters Xevo TQ-MS in positive electrospray ionization mode with multiple reaction mode was employed. Quantification of all metabolites was performed using MassLynx V4.1 software from Waters. The seven point linear calibration curves with internal standardization and 1/x weighing was constructed for the quantification.

### ChIP-Seq sample preparation

Three 15 cm dishes with cells at 70-80 % of confluency were used for one ChIP experiment. Briefly, cells were cross-linked with 1% formaldehyde for 10 minutes at room temperature, and then quenched with 125 mM glycine for 5 minutes at room temperature. Then, cells were washed with cold PBS, collected in 15 ml tubes and washed again with cold PBS by centrifugation at 1200 rpm for 5 minutes at 4 °C and finally snap frozen.

ChIP was performed as described ^41^ by using BRD4 (Bethyl Laboratories, Inc.) and MTHFD1 (sc-271413, Santa Cruz) antibodies. In brief, crosslinked cell lysates were sonicated in order to shred the chromatin into 200-500 bp fragments. Fragmented chromatin was incubated overnight at 4 °C with antibodies, followed by 2 hours at 4 °C with pre-blocked Dynabeads Protein G (ThermoFisher Scientific). Beads were washed twice with low salt buffer, twice with high salt buffer, twice with LiCl buffer, twice with 1xTE buffer and finally eluted with elution buffer for 20 min at 65 °C. The elution products were treated with RNaseA for 30 minutes at 37 °C, followed by proteinase K treatment at 55 °C for 1 hour, and then incubated at 65 °C overnight to reverse the crosslinks. The samples were further purified by using a PCR purification kit (Qiagen). ChIP-seq libraries were sequenced by the Biomedical Sequencing Facility at CeMM using the Illumina HiSeq3000/4000 platform and the 50-bp single-end configuration.

### ChIP-seq data analysis

Reads containing adapters were trimmed using Skewer ^42^ and aligned to the hg19/GRCh37 assembly of the Human genome using Bowtie2 ^43^ with the “--very-sensitive” parameter and duplicate reads were marked and removed with sambamba. Library quality was assessed with the phantomPeakQualtools scripts ^44^. For visualization exclusively, we generated genome browser tracks with the genomeCoverageBed command in BEDTools ^45^ and normalized such that each value represents the read count per base pair per million mapped and filtered reads. This was done for each sample individually and for replicates merged. In visualizations, we simply subtracted the respective merged control IgG tracks from each merged IP using IGV ^46^. We used HOMER findPeaks ^47^ in “factor” mode to call peaks on both replicates with matched IgG controls as background and used DiffBind ^48^ to detect differential binding of BRD4 or MTHFD1 in H3K27ac peaks dependent on dBET6 treatment. The top 500 differential regions for each comparison (sorted by p-value) were used for visualization using SeqPlots ^49^ and clustering with using the concentration values of each factor in each condition estimated with DiffBind. The same top differential regions were input into Enrichr ^50^ as BED files and enrichments for Reactome pathway were retrieved.

### Mouse xenograft studies

Mouse xenograft studies were performed as described previously ^25^. 2×10^6^ A549 cells, diluted 1:1 in matrigel, were transplanted subcutaneously into NOD SCID gamma mice. Treatment (30 mg/kg (*S*)-JQ1 by intraperitoneal injection five times per week, and 25 mg/kg MTX per intraperitoneal injection twice weekly) was started when tumors were established, 19 days post transplantation. Tumor volumes were evaluated twice a week by measuring two perpendicular diameters with calipers. Tumor volume was calculated using the following equation: (width*width*length)/2. Treatment was performed according to an animal licence protocol approved by the Bundesministerium für Wissenschaft und Forschung (BMWF-66.009/0280-II/3b/2012). At day 43 mice were sacrificed and tumors excised and weighted.

## ACKNOWLEDGEMENTS

S.S. is a JDRF postdoctoral fellow (3-PDF-2014-206-A-N). D.L.B. is a Merck Fellow of the Damon Runyon Cancer Research Foundation (DRG-2196-14). Next generation sequencing was performed by the Biomedical Sequencing Facility at CeMM. Research in the Kubicek lab is supported by the Austrian Federal Ministry of Science, Research and Economy, the National Foundation for Research, Technology, and Development. We thank all the members of the BioOptic Facility of the Research Institute of Molecular Pathology (IMP) and the Institute of Molecular Biotechnology GmbH (IMBA) for their help with cell sorting, Patrick Stover (Cornell) and Sebastian Nijman (Oxford) for kindly providing plasmids.

## AUTHOR CONTRIBUTIONS

S.S. and S.K. conceived the project and designed the study; S.S. and G.H. performed the gene-trap screen. F.S. and R.K. analyzed gene-trap screening data; P.R., O.H., R.I., K.M., and J.Z. performed and analyzed the BRD4 interactome screen; S.S., G.H., A.Ri.,W.Y., and S.Sch. performed biochemical and cell biological experiments and generated ChIP and metabolomics samples; K.K, B.G. and A.M. performed and analyzed proteomics and metabolomics experiments. A.Re., M.O., C.S, M.F., M.S., T.P., and C.B. performed next-generation sequencing and analyzed ChIP data; F.K. performed molecular modeling; P.M., M.O., K.P., and K.L.B. generated and analyzed chromatin-bound proteomes; P.B. and J.M. analyzed metabolomics data; D.L.B., J.E.B., and G.E.W. designed, synthesized and provided degronimids; H.P.M. and E.C. designed and conducted mouse xenograft studies. S.S. and S.K. wrote the manuscript with input from all co-authors.

## COMPETING INTERESTS

S.S. and S.K. have filed a patent application based on findings described in this manuscript.

## EXTENDED DATA

**Extended Data Figure 1.**
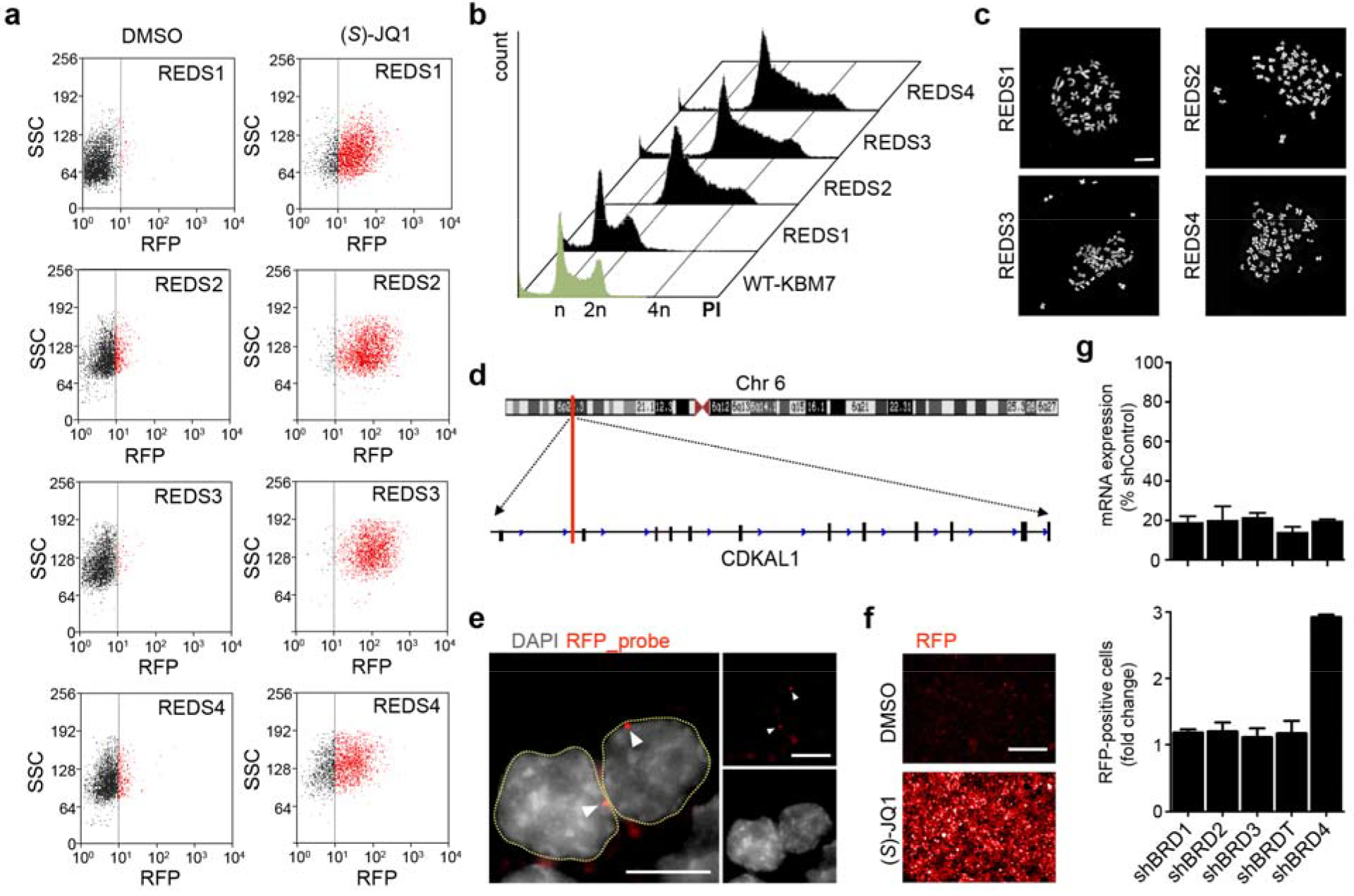
Characterization and validation of REDS BRD4 reporter clones. **a**, Representative FACS panels showing REDS1, REDS2, REDS3 and REDS4 cells treated with 0.5 μM (*S*)-JQ1; an equal volume of DMSO was used as control. Shown are representative data from three biological replicates for each experimental condition. **b**, Representative cell cycle profiles evaluated by PI-staining and DNA content analysis by FACS. REDS1, REDS3 and REDS4 cells are compared to haploid WT-KBM7 (green profile). Three biological replicates were done for each experimental condition. **c**, Metaphase chromosome spreads in REDS clones. Nuclei were stained with DAPI; scale bar 10 μm. Three biological replicates were done for each experimental condition. **d**, Representation of the RFP integration site. RFP (red bar) is inserted in the antisense direction at chromosome 6 (chr6:20,520,542-20,588,419), in the first intron of the CDKAL1 gene (sense direction). **e**, Representative fluorescence *in situ* hybridization (FISH) images in REDS1 cells. RFP probe (red dots) stains the RFP insertion. DAPI (grey signal) was used to stain the nucleus. Yellow dashed lines mark nuclear perimeter. Scale bar 10 μm. **f**, Representative live cell images of REDS1 treated with 1 μM (*S*)-JQ1 for 24 hours; an equal volume of DMSO was used as control. RFP expression is shown in red; Scale bar 100 μm. **g**, Upper panel: BRD1, BRD2, BRD3, BRDT and BRD4 expression assessed by RT-PCR following shRNA-mediated downregulation of the respective protein in REDS1 cells. Three biological replicates were done for each experimental condition (mean±STD). Lower panel: Quantification of RFP positive cells from live images of BRD1, BRD2, BRD3, BRDT or BRD4 downregulated REDS1 cells. Three biological replicates were done for each experimental condition (mean±STD).

**Extended Data Figure 2.**
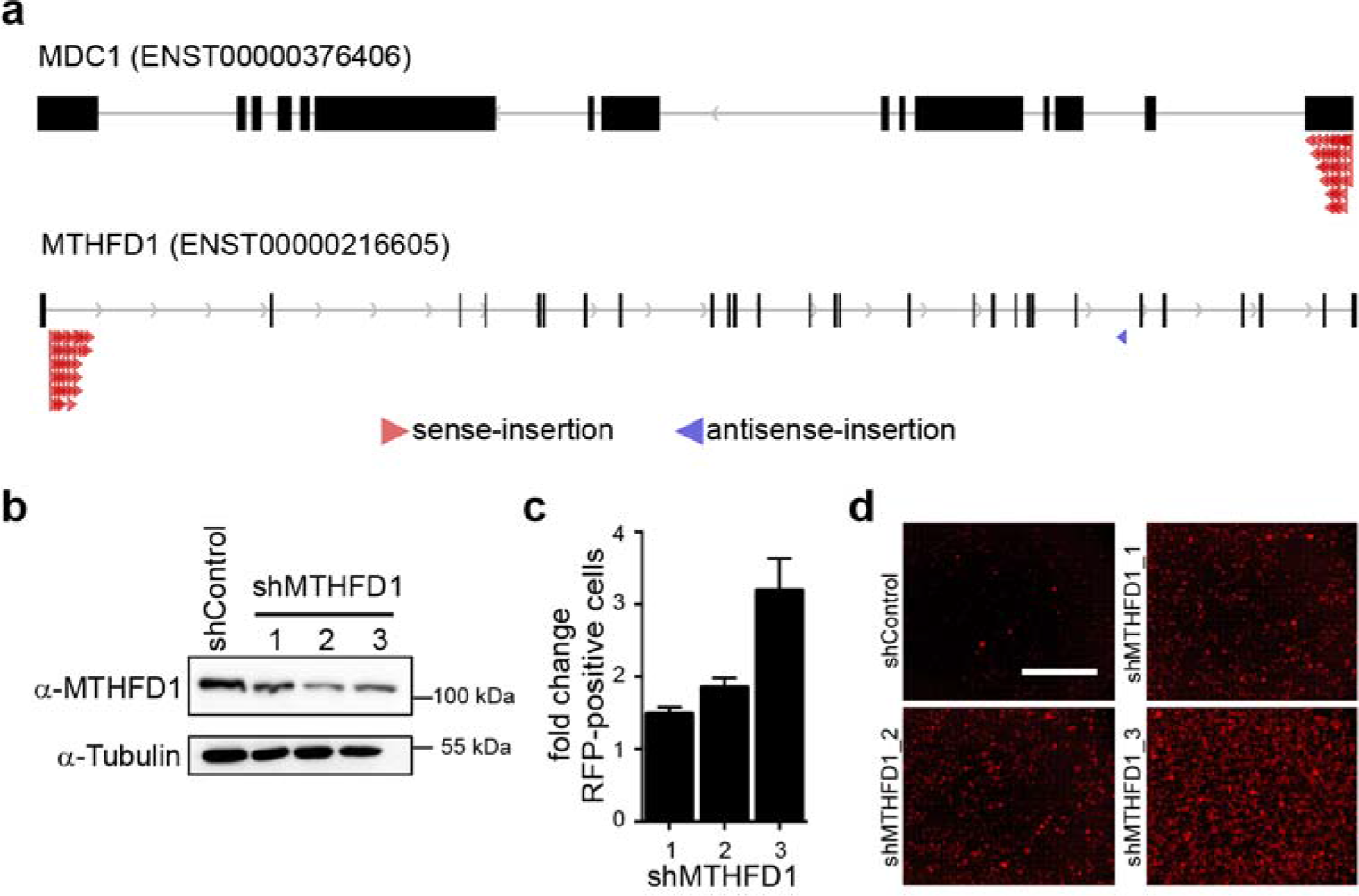
Mapping of gene trap integrations and validation of MTHFD1. **a**, Representation of the gene-trap integration sites mapped on MDC1 and MTHFD1 genes. Red arrows indicate sense insertions; blue arrows indicate antisense insertions. **b**, Western Blot for MTHFD1 protein levels after downregulation with the indicated shRNAs in REDS3. Tubulin was used as loading control. **c**, Validation of MTHFD1 in an alternative REDS clone described previously ^17^. Quantification of RFP positive cells from live-cell imaging pictures of REDS3 cells treated with MTHFD1 shRNA. Three biological replicates were done for each experimental condition (mean±STD). **d**, Representative live-cell images of MTHFD1 knock down REDS3 cells. RFP signal is shown in red; scale bar 100 μm.

**Extended Data Figure 3.**
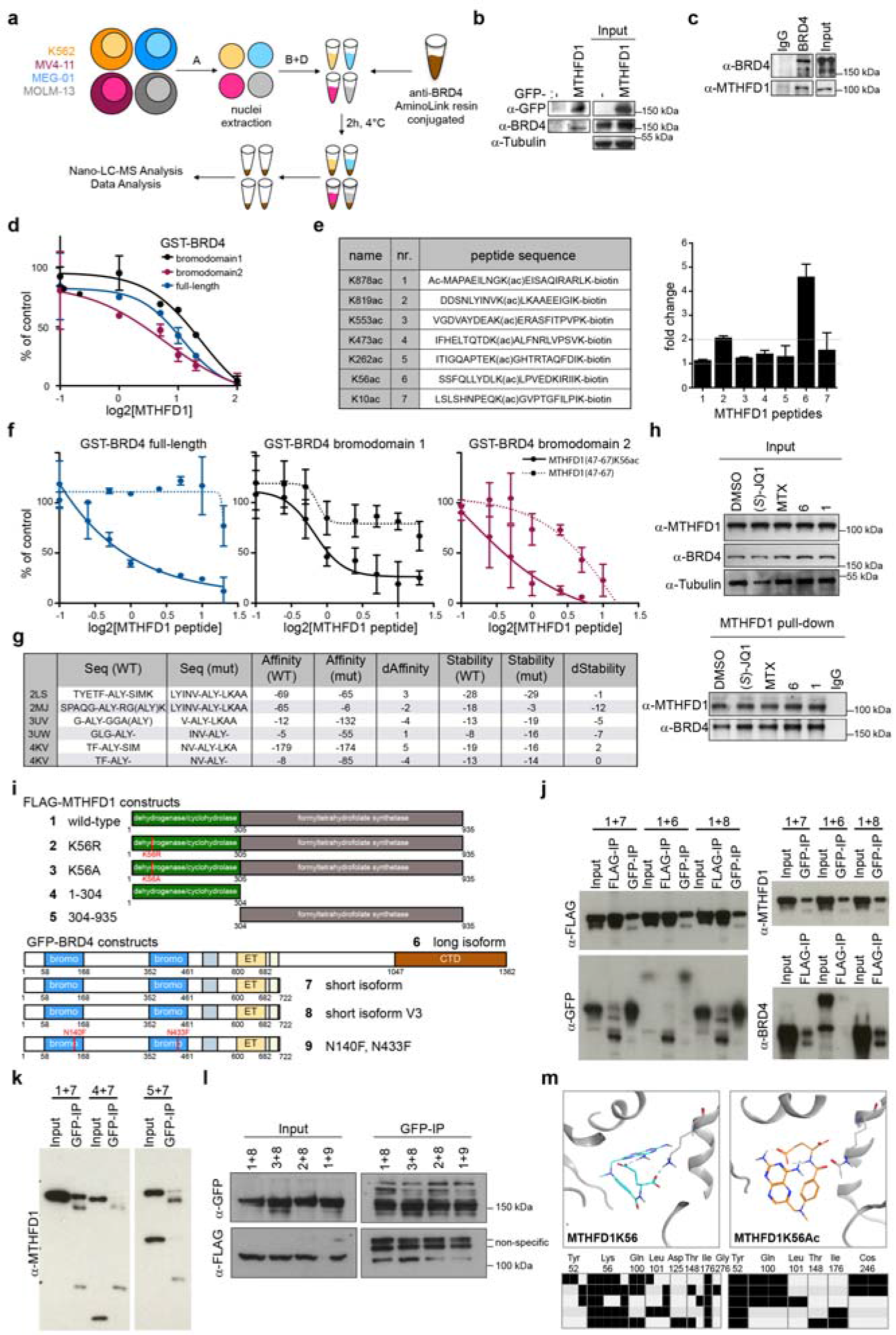
Mapping the BRD4-MTHFD1 interaction site. **a**, Experimental setup of BRD4 interaction proteomics. **b**, Western blot showing GFP pull-down from HEK293T whole-cell extracts overexpressing GFP-MTHFD1 or GFP alone. Tubulin was used as loading control. **c**, Western blot showing BRD4 pull-down from HeLa whole-cell extracts. **d**, AlphaLISA for the interaction between BRD4 and an acetylated histone peptide in the presence of increasing amounts of recombinant MTHFD1. **e**, AlphaLISA assay with the indicated MTHFD1 acetylated peptides and GST-BRD4 (full length). Two replicates for each condition (mean±STD). **f**, Titration of MTHFD1 peptides unmodified and K56 acetylated for their ability to disrupt the interaction of GST-BRD4 with an acetylated histone peptide. Two replicates for each condition (mean±STD). **g**, Predicted affinity of BRD4 to the MTHFD1(K56ac) peptide (Affinity(mut)) compared to the histone peptide co-crystalized (Affinity(WT)). Negative scores are indicative of higher affinity. Similarly, stabilities of the peptide in the bound configuration calculated for the original histone (Stability(WT)) and the MTHFD1(K56ac) peptides (Stability(mut)). **h**, MTHFD1 pull-down on HAP1 whole-cell lysates treated with 50 μM (S)-JQ1, MTX, peptides MTHFD1K56Ac (6) or MTHFD1K878Ac (1; negative control). An equal amount of DMSO was used as control, tubulin was used as loading control. **i**, MTHFD1 and BRD4 constructs used in immunoprecipitation experiments shown in panels J-L. **j**, FLAG and GFP immunoprecipitation from HEK293 cells overexpressing the indicated constructs shows interaction of MTHFD1 with all three BRD4 isoforms. **k**, GFP immunoprecipitation from HEK293 cells overexpressing the indicated constructs shows interaction of BRD4 with full-length MTHFD1 but not with the individual domains of the protein. **l**, GFP immunoprecipitation from HEK293 cells overexpressing the indicated constructs shows increased interaction of MTHFD1(K56A) and decreased interaction of MTHFD1(K56R) with the short isoform of BRD4. Similarly, the interaction is impaired in the BRD4 double bromodomain mutant. **m**, Docking of Methotrexate to the binding pocked of MTHFD1. Methotrexate is predicted to interact with Lysine 56 of MTHFD1 (left panel), and this interaction is lost when K56 is acetylated (right panel). Protein Ligand Interaction Fingerprints (PLIF) for all docking poses are shown at the bottom.

**Extended Data Figure 4.**
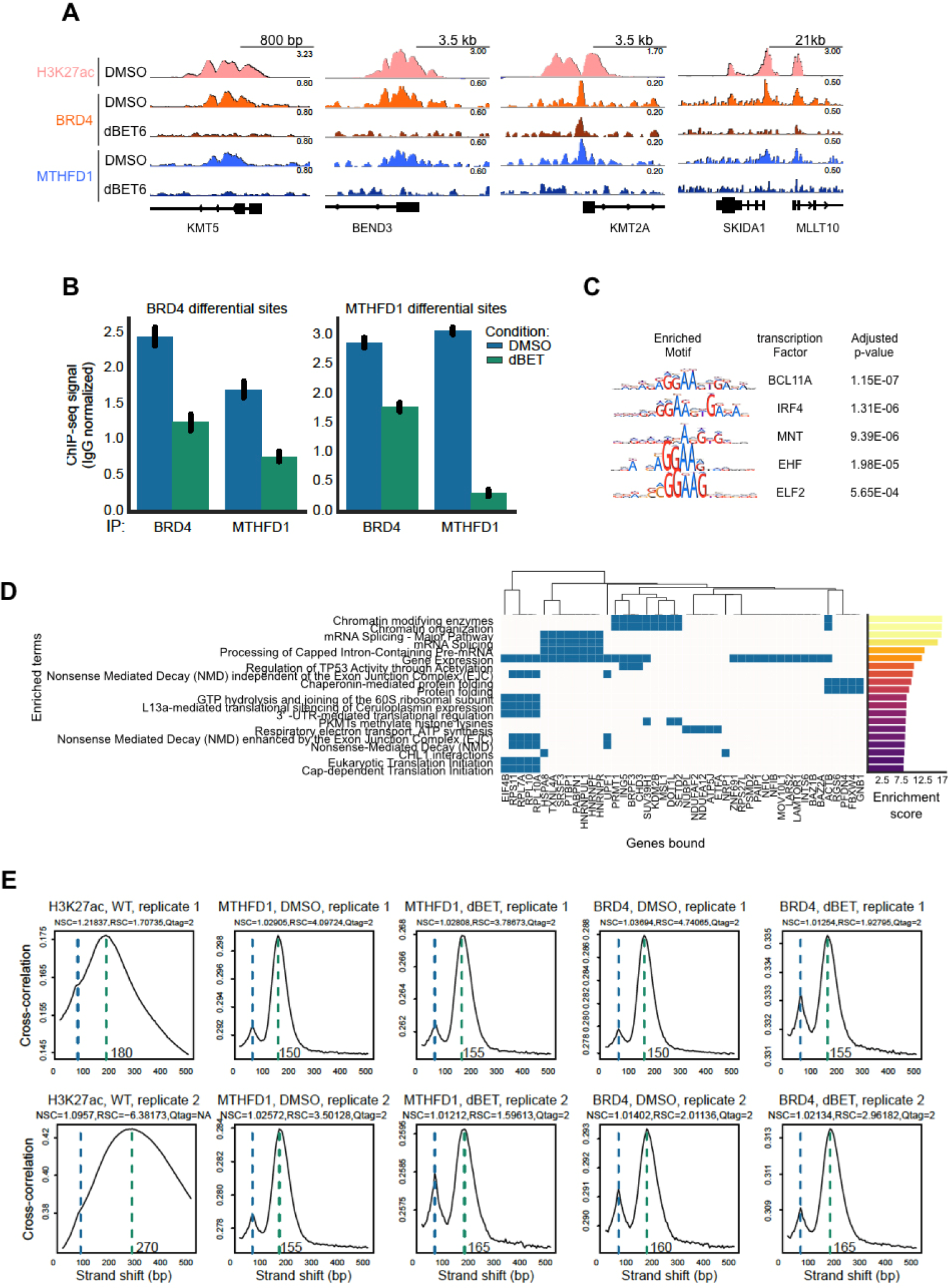
Locus-specific binding of MTHFD1. **a**, Illustrative genome browser views of BRD4 and MTHFD1 binding in the H3K27ac-marked promoters of KMT5A, BEND3, KMT2A, SKIDA1 (from left to right). All tracks were normalized by the total mapped reads in the genome and the respective IgG control was subtracted from the merged replicate tracks. Tracks for the same factor in different conditions were scaled similarly for comparison. Note the loss of BRD4 and MTHFD1 binding upon 1 μM dBET6 treatment for 2 hours. **b**, Quantification of BRD4 and MTHFD1 in the top 500 differentially bound sites by MTHFD1 or BRD4 between dBET6 or DMSO treatment. Values represent estimated factor abundance normalized by matched IgG signal. Error bars represent 95th confidence interval. **c**, Enriched motifs found in the joint set of regions with differential BRD4 or MTHFD1 binding upon dBET6 treatment. Note the recurrent “GGAA” motif found. **d**, Reactome pathway terms enriched in genes bound differentially by BRD4 or MTHFD1 binding upon dBET6 treatment. The central heatmap illustrates which genes belong to each term. The enrichment score on the right represents “log(p-value) * Z-score”, where the Z-score is the gene set’s deviation from an expected rank as defined by Enrichr. **e**, Quality of ChIP-seq libraries through cross-correlation analysis. The X-axis represents an amount (in base pairs) by which the signal in two aligned strands is shifted by, and the Y-axis represents the cross-correlation between the signal in the strands at each shifted position. The first increase in cross-correlation (marked by a blue dashed line) is noise related with the read length used, and the second is true signal (green dashed line) related with enrichment of the immunoprecipitated protein and generally reflects the average length of DNA bound by the protein. The amount of baseline-normalized cross-correlation (NSC) and ratio between the two cross-correlation values (RSC) is indicative of signal-to-noise ratio and therefore of library quality (Qtag, increasing from 0 to 2).

**Extended Data Figure 5.**
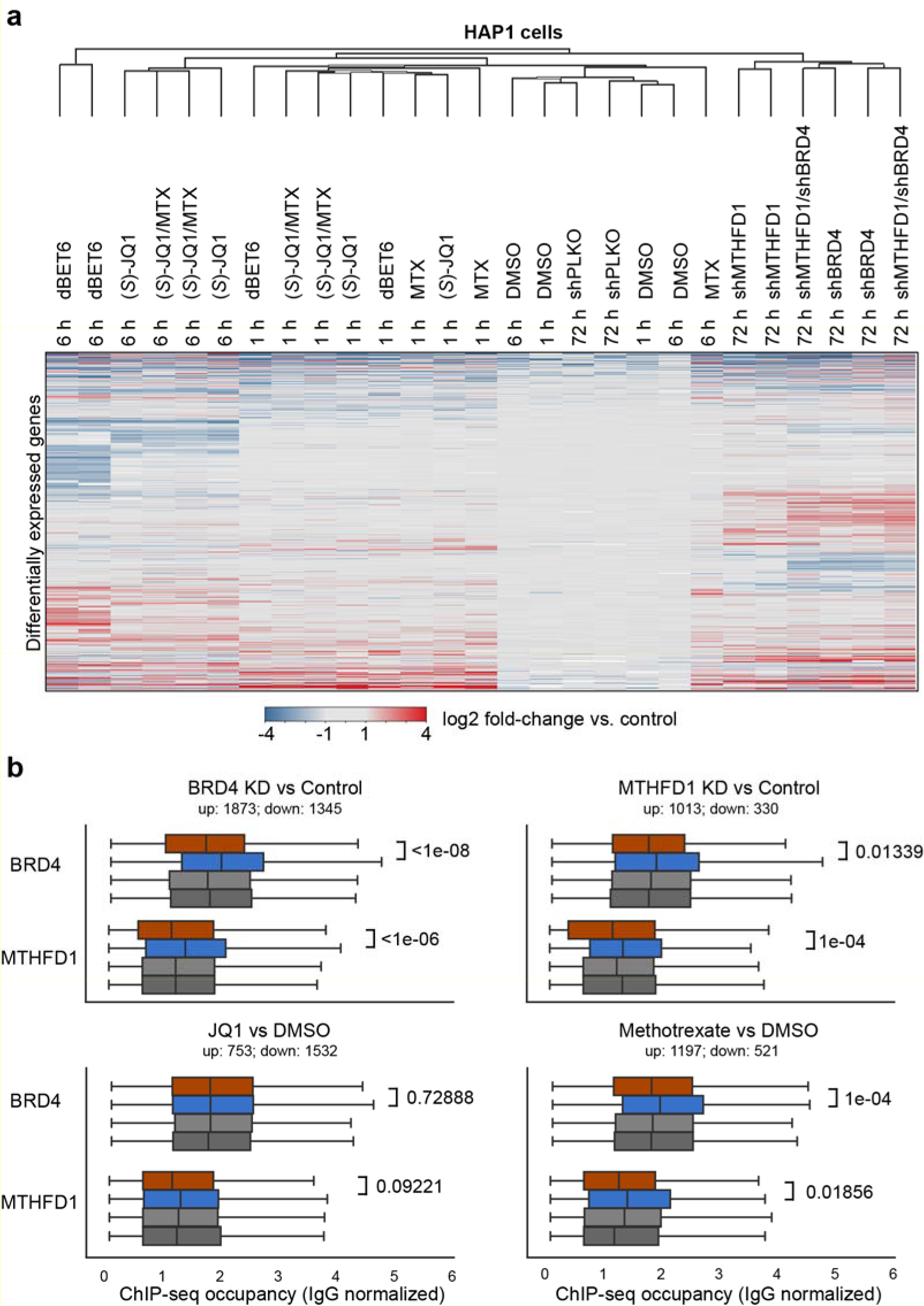
Transcriptional consequences of targeting MTHFD1 and BRD4 in HAP1 cells. **a**, Heat map of relative transcriptional changes of HAP1 cells treated with 0.1μM dBET6, 1μM (*S*)-JQ1, 1μM MTX, shRNAs targeting BRD4 or MTHFD1 alone or in combination. Equal amount of DMSO, or non-targeting hairpins were used as respective control conditions. **b**, Integration of ChIP-Seq and RNA-Seq data in HAP1 cells. BRD4 and MTHFD1binding at sites associated with genes which are up- (orange) or down-regulated (blue) upon knockdown of either BRD4, MTHFD1, or treatment with either JQ1 or Methotrexate. Binding in random sets of genes of the same size as the respective up- or down-regulated sets is displayed as control. Values represent estimated factor abundance normalized by matched IgG signal and equality of distributions was assessed with the Kolmogorov-Smirnov test.

**Extended Data Figure 6.**
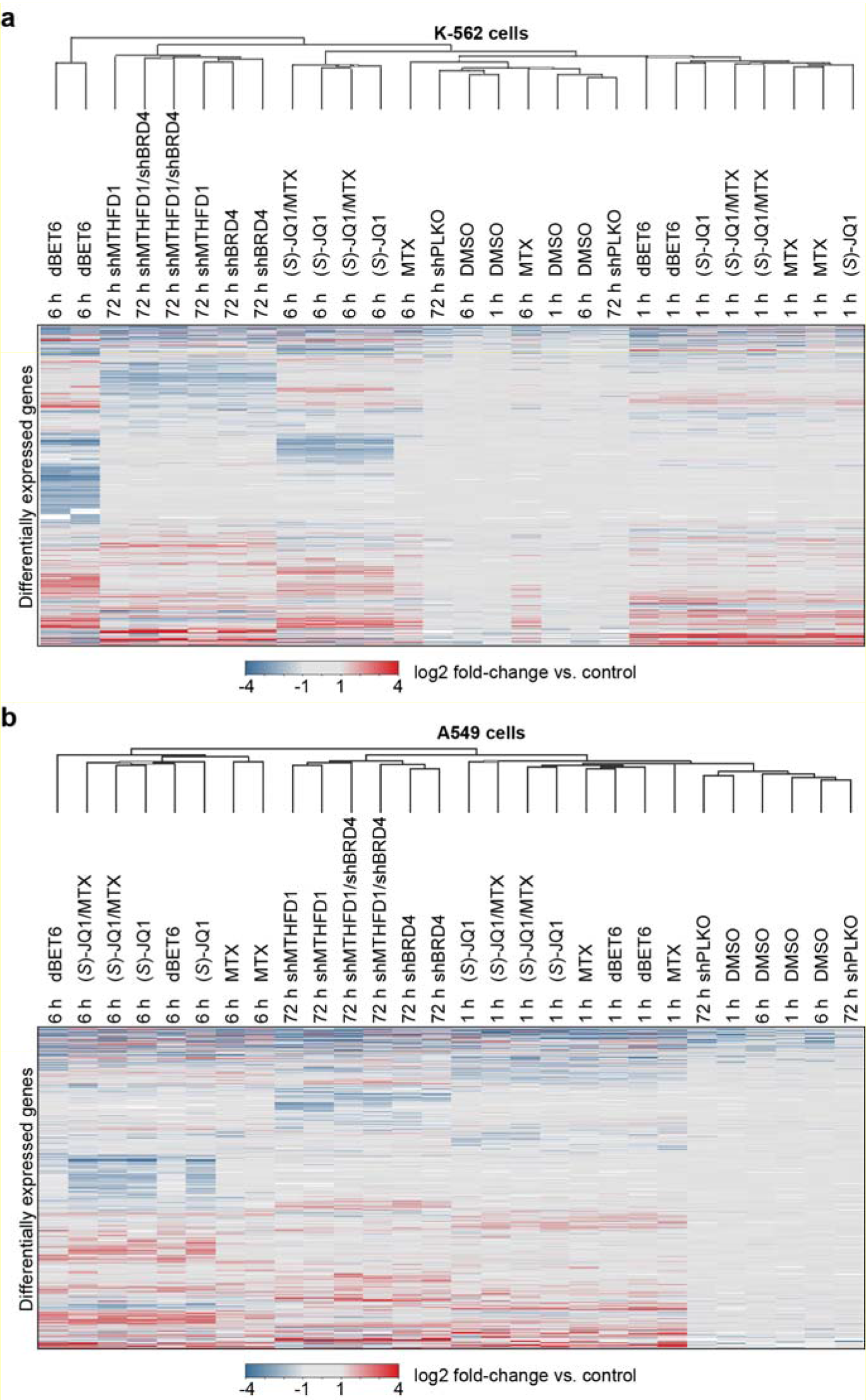
Transcription changes in K-562 and A549 cells. **a**, Heat map of relative transcription changes in K-562 cells treated with with 0.1μM dBET6, 1μM (*S*)-JQ1, 1μM MTX, shRNAs targeting BRD4 or MTHFD1 alone or in combination. Equal amount of DMSO, or non-targeting hairpins were used as respective controls. **b**, Heat map matrix of relative transcription changes in A549 treated with 0.1μM dBET6, 1μM (*S*)-JQ1, 1μM MTX, shRNAs targeting BRD4 or MTHFD1 alone or in combination. Equal amount of DMSO, or non-targeting hairpins were used as respective controls.

**Extended Data Figure 7.**
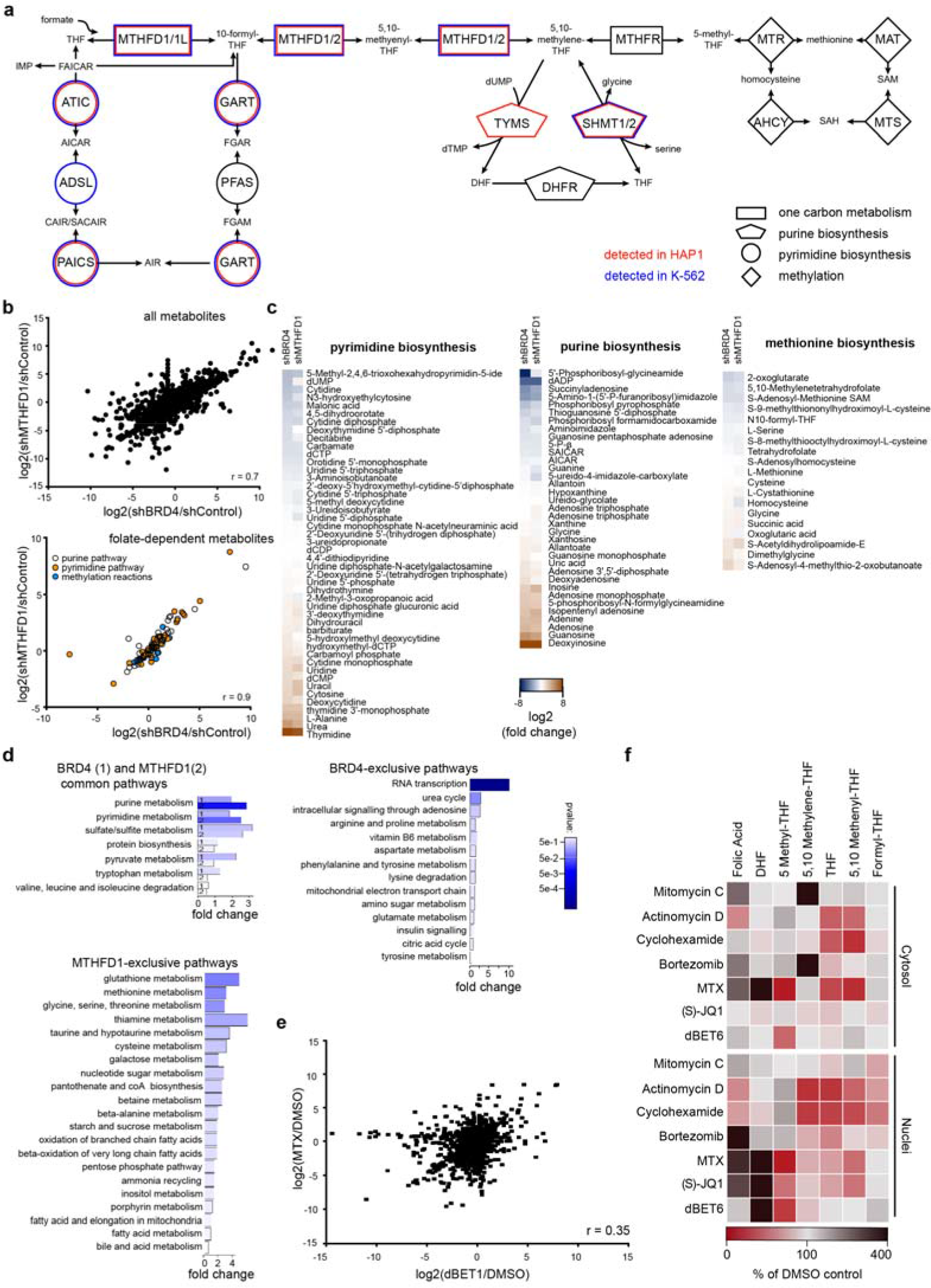
Effects of MTHFD1 loss on nuclear metabolite composition. **a**, Representation of the folate pathway. Enzyme names are reported inside the geometric shapes, connecting the different metabolites. Enzymes that were found associated with chromatin in HAP1 and K-562 cells by mass spectrometry analysis are indicated in red and blue, respectively. Two biological replicates were done. **b**, Scatter plot depicting changes in nuclear metabolite levels following BRD4 or MTHFD1 downregulation for all metabolites (upper panel) and folate-dependent metabolites (lower panel). Means from two biological replicates. **c**, Heat maps representing metabolite changes in the pyrimidine, purine and methionine biosynthetic pathways upon downregulation of BRD4 or MTHFD1 by shRNA. Two biological replicates were done for each experimental condition. **d**, Metabolite Set Enrichment Analysis (MSEA) showing a comparison of the enrichment of metabolite sets that are significant in both knockdowns (shBRD4: 1; shMTHFD1: 2), metabolite sets exclusively enriched in shMTHFD1 or metabolite sets exclusively enriched in shBRD4. **e**, Correlation plot depicting changes in nuclear metabolite levels following dBET1 or MTX treatment of HAP1 cells for 24 h. Means from two biological replicates. **f**, Heatmaps showing relative changes in folate metabolites levels in the of nuclear and cytosolic fraction of HAP1 cells treated with 1 μM of Mitomycin C, Actinomycin D, Bortexomib, MTX and (*S*)-JQ1, 0.5 μM of dBET6 or 12.5 μM of Cyclohexamide for 6 hours. Equal amount of DMSO was used as control, 2 biological replicates were done for each experimental condition.

**Extended Data Figure 8.**
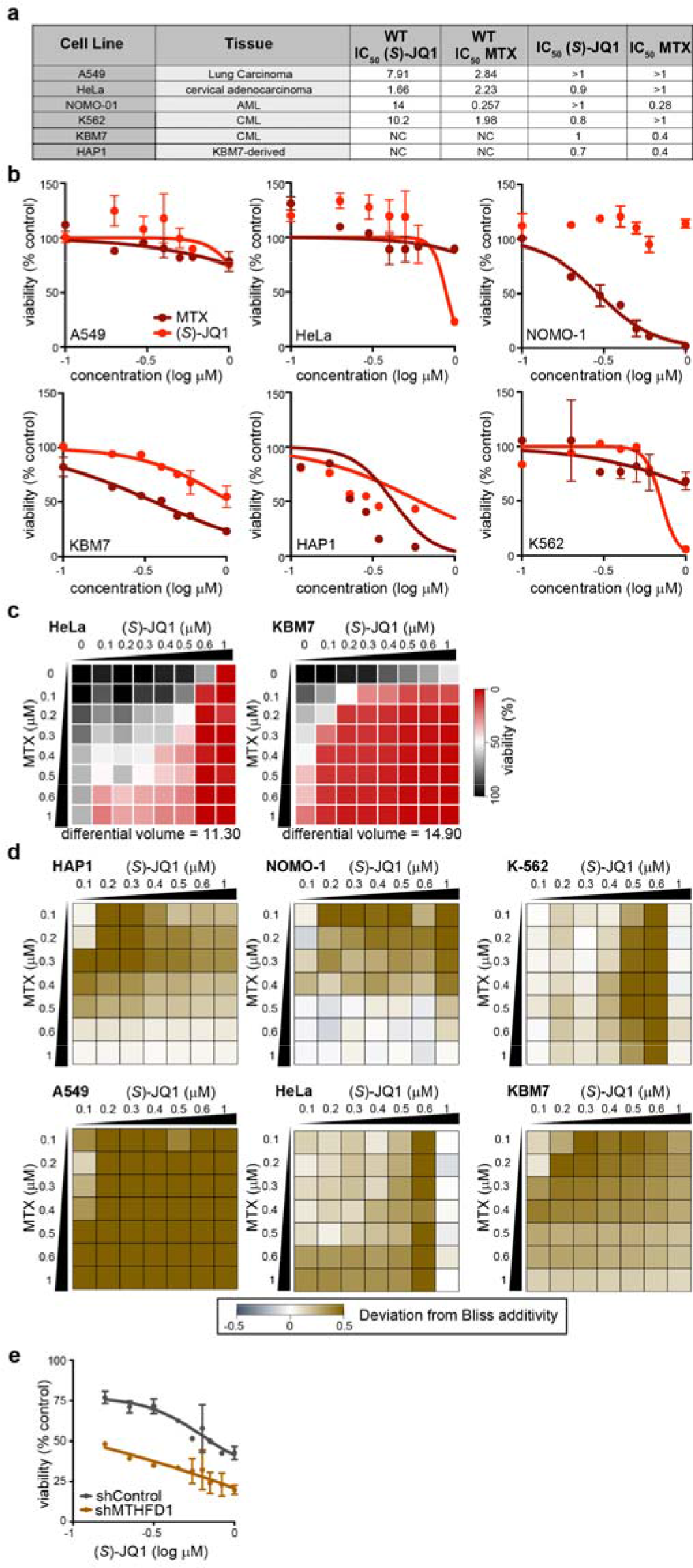
Synergism of MTX and (*S*)-JQ1. **a**, Table of IC_50_ values for (*S*)-JQ1 and MTX as reported in the Welcome Trust-Genomics of Drug Sensitivity in Cancer database (WT; http://www.cancerrxgene.org), or produced in house (right two columns), for the different cell lines. **b**, Determination of IC_50_ values for (*S*)-JQ1 (red) and MTX (dark red) in the indicated cell lines. Three biological replicates were done for each experimental condition (mean±STD). **c**, Dose response matrices displaying cell viability of HeLa and KBM7 treated for 72 h with (*S*)-JQ1 and MTX alone or in combination. Means from two biological replicates. **d**, Deviations from Bliss additivity for drug matrices from Fig. 4C and S6C. **e**, Knock-down of MTHFD1 in A549 cells followed by 72 h treatment with increasing concentrations of (*S*)-JQ1.

**Extended Data Table 1.**
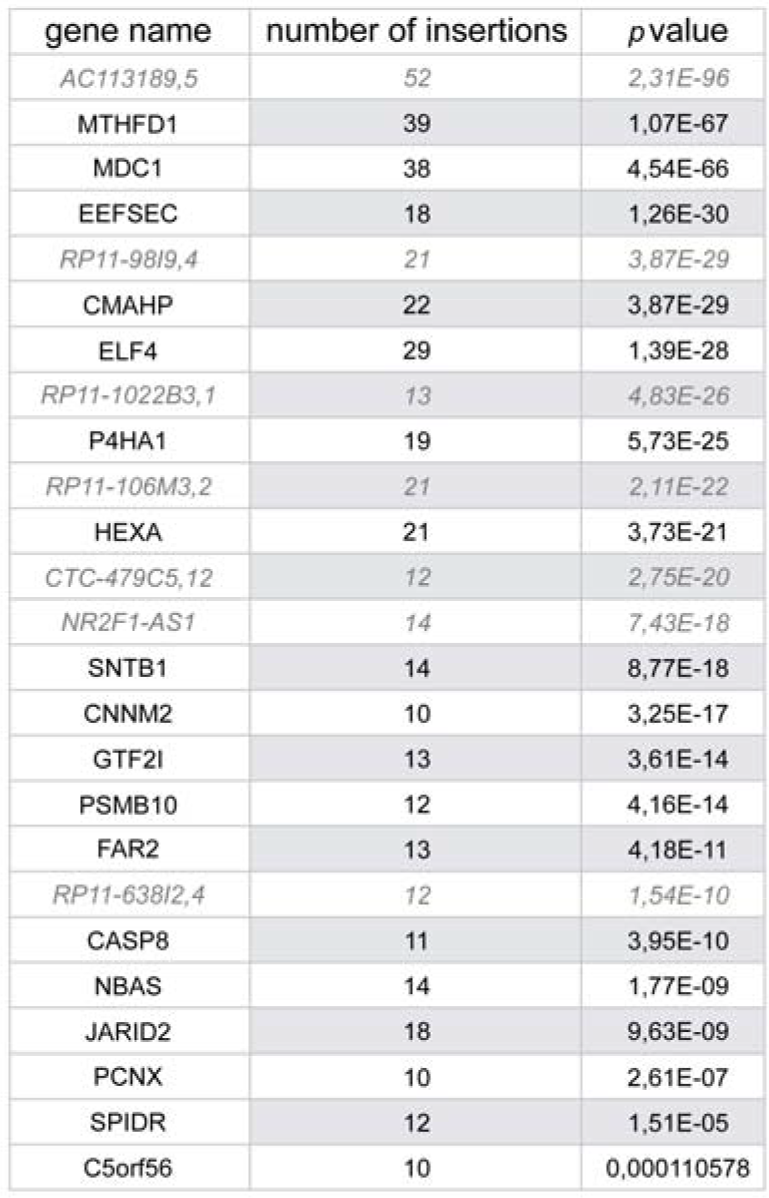
List of the significant gene-trapped loci. The loci reported in the table were selected if showing more than 10 insertions in combination with a significant *p-*value. In grey and italic are the identified long non-coding RNAs (not reported in the circus plot of Fig. 1C); in black and regular the identified coding genes. The values were calculated using the Fisher test, as described in the methods section.

**Extended Data Table 2.**
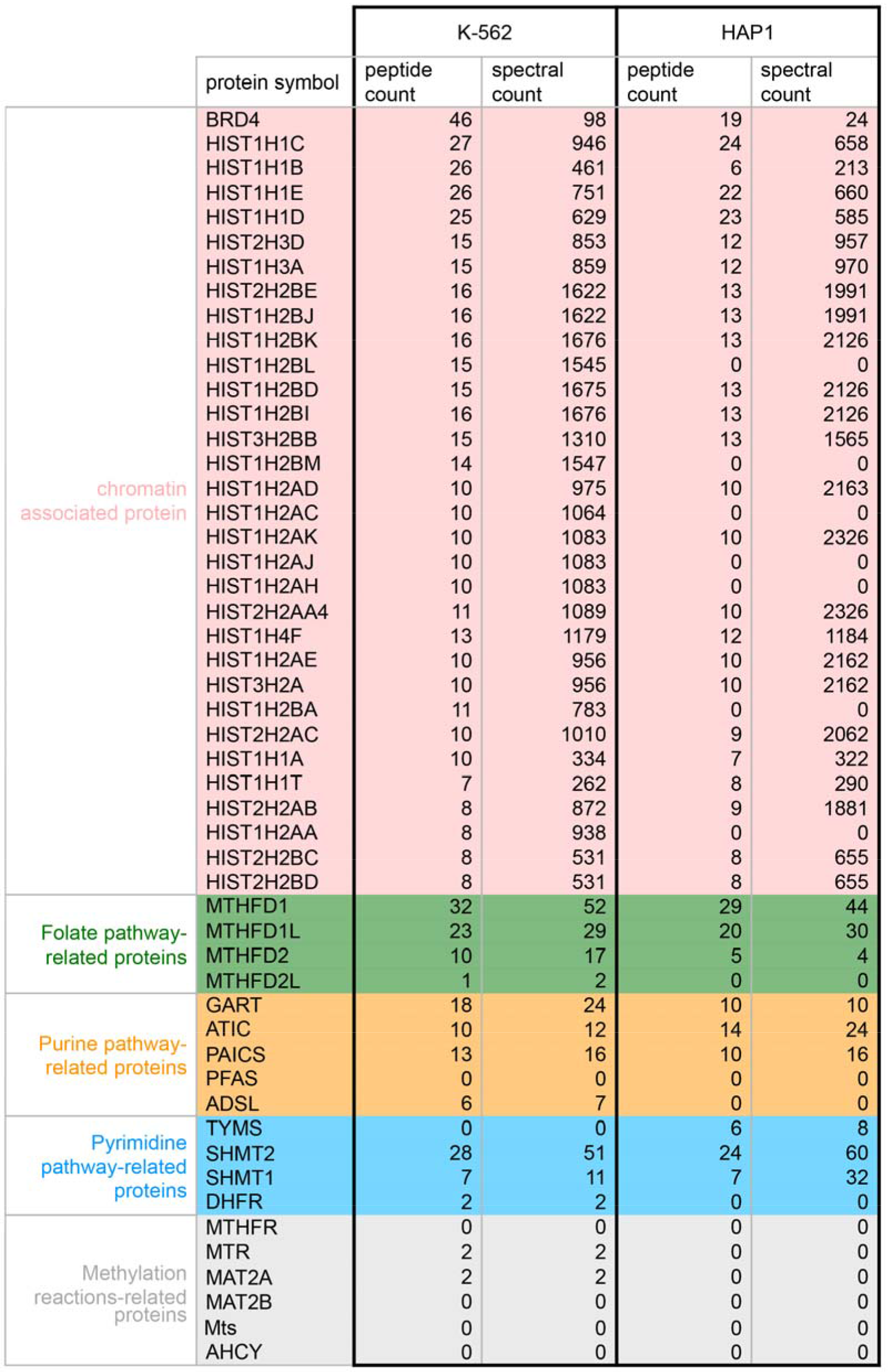
List of the chromatin-associated folate-pathway enzymes. Peptide and spectral counts identified by MS analysis of HAP1 chromatin extracts. Two biological replicates were done. Enzymes of the folate pathway found associated with chromatin are written in regular, while in italic are the “not-found”. BRD4 and histones are shown as control for chromatin associated proteins.

|  | protein symbol | K-562 |  | HAP1 |  |
| --- | --- | --- | --- | --- | --- |
|  |  | peptide count | spectral count | peptide count | spectral count |
| chromatin associated protein | BRD4 | 46 | 98 | 19 | 24 |
|  | HIST1H1C | 27 | 946 | 24 | 658 |
|  | HIST1H1B | 26 | 461 | 6 | 213 |
|  | HIST1H1E | 26 | 751 | 22 | 660 |
|  | HIST1H1D | 25 | 629 | 23 | 585 |
|  | HIST2H3D | 15 | 853 | 12 | 957 |
|  | HIST1H3A | 15 | 859 | 12 | 970 |
|  | HIST2H2BE | 16 | 1622 | 13 | 1991 |
|  | HIST1H2BJ | 16 | 1622 | 13 | 1991 |
|  | HIST1H2BK | 16 | 1676 | 13 | 2126 |
|  | HIST1H2BL | 15 | 1545 | 0 | 0 |
|  | HIST1H2BD | 15 | 1675 | 13 | 2126 |
|  | HIST1H2BI | 16 | 1676 | 13 | 2126 |
|  | HIST3H2BB | 15 | 1310 | 13 | 1565 |
|  | HIST1H2BM | 14 | 1547 | 0 | 0 |
|  | HIST1H2AD | 10 | 975 | 10 | 2163 |
|  | HIST1H2AC | 10 | 1064 | 0 | 0 |
|  | HIST1H2AK | 10 | 1083 | 10 | 2326 |
|  | HIST1H2AJ | 10 | 1083 | 0 | 0 |
|  | HIST1H2AH | 10 | 1083 | 0 | 0 |
|  | HIST2H2AA4 | 11 | 1089 | 10 | 2326 |
|  | HIST1H4F | 13 | 1179 | 12 | 1184 |
|  | HIST1H2AE | 10 | 956 | 10 | 2162 |
|  | HIST3H2A | 10 | 956 | 10 | 2162 |
|  | HIST1H2BA | 11 | 783 | 0 | 0 |
|  | HIST2H2AC | 10 | 1010 | 9 | 2062 |
|  | HIST1H1A | 10 | 334 | 7 | 322 |
|  | HIST1H1T | 7 | 262 | 8 | 290 |
|  | HIST2H2AB | 8 | 872 | 9 | 1881 |
|  | HIST1H2AA | 8 | 938 | 0 | 0 |
|  | HIST2H2BC | 8 | 531 | 8 | 655 |
|  | HIST2H2BD | 8 | 531 | 8 | 655 |
| Folate pathway-related proteins | MTHFD1 | 32 | 52 | 29 | 44 |
|  | MTHFD1L | 23 | 29 | 20 | 30 |
|  | MTHFD2 | 10 | 17 | 5 | 4 |
|  | MTHFD2L | 1 | 2 | 0 | 0 |
| Purine pathway-related proteins | GART | 18 | 24 | 10 | 10 |
|  | ATIC | 10 | 12 | 14 | 24 |
|  | PAICS | 13 | 16 | 10 | 16 |
|  | PFAS | 0 | 0 | 0 | 0 |
|  | ADSL | 6 | 7 | 0 | 0 |
| Pyrimidine pathway-related proteins | TYMS | 0 | 0 | 6 | 8 |
|  | SHMT2 | 28 | 51 | 24 | 60 |
|  | SHMT1 | 7 | 11 | 7 | 32 |
|  | DHFR | 2 | 2 | 0 | 0 |
| Methylation reactions-related proteins | MTHFR | 0 | 0 | 0 | 0 |
|  | MTR | 2 | 2 | 0 | 0 |
|  | MAT2A | 2 | 2 | 0 | 0 |
|  | MAT2B | 0 | 0 | 0 | 0 |
|  | Mts | 0 | 0 | 0 | 0 |
|  | AHCY | 0 | 0 | 0 | 0 |

